# Nucleoside Binding by a Surface Lipoprotein Governs Conjugative ICE Acquisition in Ruminant Mycoplasmas

**DOI:** 10.1101/2025.09.22.677790

**Authors:** M’hamed Derriche, Laurent Xavier Nouvel, Calvin Fauvet, Núria Mach, Elisa Simon, Gwendoline Pot, Hortensia Robert, Alexandre Stella, Christian de la Fe, Renaud Maillard, Sergi Torres-Puig, Yonathan Arfi, Christine Citti, Eric Baranowski

**Author notes:** Address correspondence to Eric Baranowski.

## Abstract

Integrative and conjugative elements (ICEs) are major mediators of horizontal gene transfer (HGT) in bacteria. However, the role of recipient cells in their acquisition has received little attention. Using the ruminant pathogens *Mycoplasma agalactiae* and *Mycoplasma bovis* as minimal models, we combined genome-wide transposon mutagenesis with high-throughput mating assays to identify recipient factors required for ICE acquisition. The surface lipoprotein P48 emerged as the primary determinant of ICE uptake in both species. Structural and functional analyses revealed that P48 is the substrate-binding component of an ABC transporter with nucleoside-binding capacity. A single point mutation that abolished nucleoside binding drastically reduced ICE acquisition, demonstrating that P48-mediated nucleoside recognition is essential for conjugative transfer. However, ICE uptake did not require nucleoside transport, as inactivation of the transporter permease blocked nucleoside analog toxicity but not ICE invasion. Loss of P48 also triggered transcriptional activation of vestigial ICE genes, suggesting that surface recognition affects the intracellular state of the recipient. Remarkably, ICE transmission from recipient-derived donors was unaffected by P48 loss, underscoring its acquisition-specific role. Together, these results reveal a previously unrecognized, surface-exposed recipient factor critical for efficient ICE transfer in mycoplasmas and identify nucleotide binding as a central function in conjugation. By demonstrating that recipient-encoded functions can directly control ICE dissemination, this work challenges the donor-centric paradigm of bacterial conjugation and suggests new strategies to restrict horizontal gene flow in pathogenic and synthetic mycoplasmas.

**IMPORTANCE:** Integrative and conjugative elements (ICEs) are mobile DNA elements that drive bacterial conjugation, a major process by which bacteria exchange genes. Although conjugation has been studied for decades, the focus has been almost exclusively on donor cells and the ICE itself, leaving the role of recipient cells largely overlooked. Using the wall-less ruminant pathogens *Mycoplasma agalactiae* and *Mycoplasma bovis* as minimal models, we discovered that a single recipient lipoprotein is required for efficient ICE uptake. Our data show that nucleoside recognition by P48, but not transport, is critical for conjugation, revealing an unexpected mechanistic link between nutrient sensing and gene acquisition. These findings shift the paradigm of conjugation from a donor-driven process to one jointly determined by donor and recipient functions. By identifying a recipient-encoded determinant of ICE transfer, this work opens new avenues to control horizontal gene flow in both pathogenic and engineered bacteria.

**Graphical abstract:** 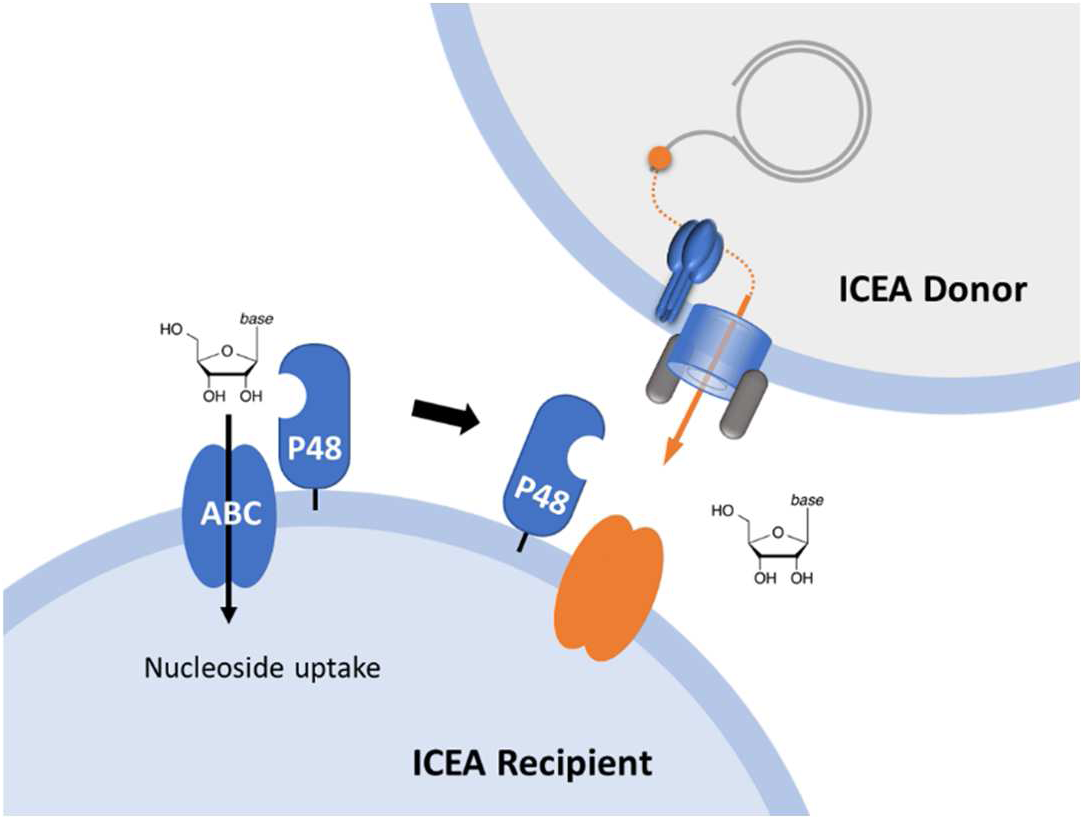

## INTRODUCTION

As major mediators of HGT, ICEs are among the most prevalent conjugative systems identified in prokaryotes, reflecting their evolutionary success and impact on microbial genome plasticity (1–6).

The recent discovery of ICEs in *Mollicutes,* commonly known as mycoplasmas, has reshaped our classical view of how these minimal bacteria evolve (7–9). Mycoplasmas originated from low G+C content Gram-positive ancestors through extensive genome reduction, which eliminated many metabolic functions and rendered them strictly dependent on host-derived nutrients (10). For decades, their evolution was regarded as degenerative, driven largely by gene loss. However, the identification of ICE-mediated horizontal gene transfer revealed that mycoplasma genomes remain dynamic and plastic, incorporating newly acquired genes in addition to losing ancestral ones (7, 9). Notably, mycoplasma ICEs (MICEs) mediate not only canonical conjugation but also mycoplasma chromosomal transfer (MCT), an unconventional mechanism in which multiple chromosomal fragments are exchanged horizontally (7, 11). MCT generates progeny with highly mosaic genomes, greatly expanding the adaptive potential of these minimal bacteria (12, 13).

The ICE of the ruminant pathogen *Mycoplasma agalactiae* (ICEA) has been the most extensively studied MICE to date (14–17). Unlike many classical ICEs that integrate at conserved chromosomal sites, ICEA inserts randomly throughout the host genome and relies on a DDE transposase of the prokaryotic Mutator-like family for mobility (15). ICEA spans 27 kb and encodes 23 genes, most without homologs outside *Mollicutes*, underscoring its unique evolutionary history (14–17). Comparative analyses have identified a minimal MICE backbone of 4 genes predicted to be essential for ICE chromosomal integration-excision and horizontal dissemination through a type IV secretion system (14, 18).

The dissemination of ICEs depends not only on their own conjugative machinery but also on host-encoded functions and environmental cues (1, 4, 19–21). In the human pathogen *Mycoplasma hominis*, stress conditions such as DNA damage can trigger ICE activation, but co-incubation with eukaryotic cells proved to be a far more potent inducer (22). This finding aligns with our recent results showing that interactions with eukaryotic cells, across various *ex vivo* infection models, markedly increase ICE transfer in ruminant mycoplasma species (23). In other bacteria, ICE activation can also be influenced by several host-encoded functions, among which the RecA-dependent SOS response is the best characterized (1, 24). Remarkably, most mycoplasmas lack a canonical SOS system, with the exception of *Mycoplasma gallisepticum,* which retains an SOS-like mechanism (25). Finally, it is worth noting that *Mycoplasma genitalium* has been shown to engage in RecA-dependent HGT independent of mobile elements (26).

While donor functions and ICE-encoded factors have been well characterized, the contribution of the recipient partner to ICE transfer, particularly during the early stages of conjugation, remains largely unexplored. Recipient defenses such as restriction–modification, CRISPR-Cas, or other barriers to foreign DNA are known to limit gene flow (27). Beyond these defenses, relatively few recipient-encoded determinants of ICE uptake have been described (28, 29), including lipoprotein maturation and cell surface architecture, which can promote or hinder donor–recipient contact, thereby influencing conjugation efficiency.

In this study, we took advantage of the streamlined genome of *M. agalactiae* and its close relative *Mycoplasma bovis* to map chromosomal regions that affect ICE acquisition by recipient cells. Using genome-wide mutagenesis and mating assays, we discovered that ICEA acquisition depends mainly on the nucleoside-binding activity of a surface-exposed lipoprotein, but nucleoside uptake itself was dispensable for ICEA acquisition. These findings challenge the traditional donor-centric view of bacterial conjugation, and identify a novel, surface-exposed function that could be exploited to limit HGT in mycoplasmas.

## RESULTS

### *M. agalactiae* genome-encoded determinants influencing ICEA acquisition

To assess the contribution of recipient functions to ICE movements in *M. agalactiae*, we mapped chromosomal regions influencing ICEA acquisition by the ICE-negative strain PG2. For this purpose, a library of 3,016 individual mutants of PG2 was generated, each carrying a stable mini-transposon (mTn) encoding a puromycin-resistance cassette (P-tag). Individual mutants were tested in 96-well plates by mating with the donor strain PG2^T^[ICEA]^G^, in which the ICEA was tagged with a gentamicin-resistance marker (G-tag) (Table 1).

**TABLE 1.**
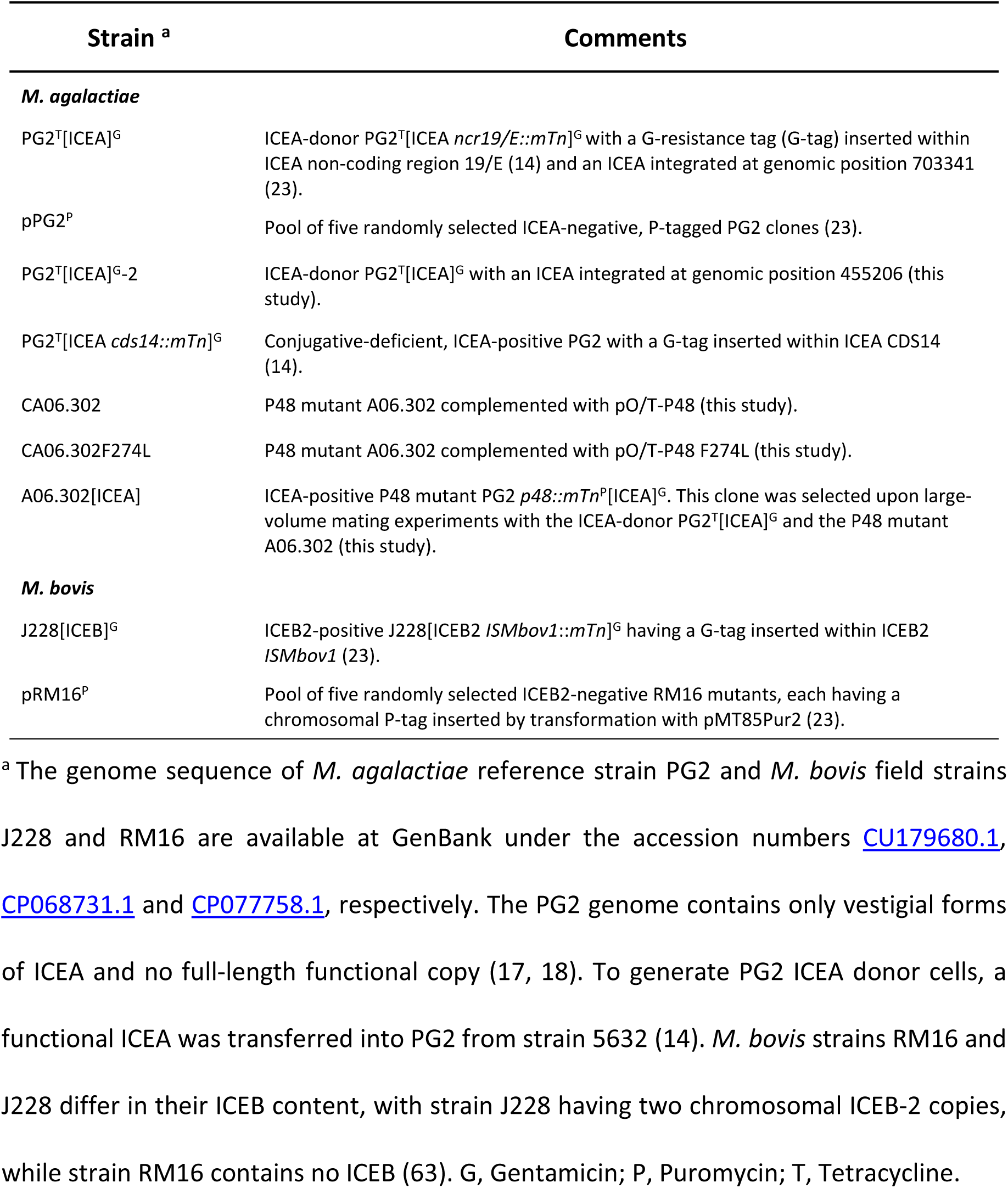
Mycoplasma strains used in mating experiments.

This high-throughput screen yielded 18 mutants resistant to ICEA invasion (Table 2). Mapping of mTn insertion sites revealed 18 unique loci distributed across 11 coding sequences (CDSs) and one non-coding region (Table 2). Eight CDSs encoded proteins with DNA-related functions, including replication (MAG4380), repair (MAG03780, MAG3790, MAG4990, MAG5020), recombination (MAG2480) and transposition (MAG3410), as well as nucleoside metabolism (MAG5120). Two others (MAG0120 and MAG0160) belonged to an ABC transporter locus, with MAG0120 encoding the predicted substrate-binding protein. One last CDS encoded a membrane protein of unknown function (MAG5810).

**Table 2.** Transposon insertion sites in selected *M. agalactiae* PG2 mutants.

| Mutant <sup>a</sup> | Genomic position <sup>b</sup> | CDS <sup>c</sup> | Relative position <sup>d</sup> | Resistance to ICEA <sup>e</sup> | KEGG orthology assignment <sup>f</sup> |
| --- | --- | --- | --- | --- | --- |
| A01.088 | 511991 (+) | MAG4380 | 0.51 (-) | + | K03529 Chromosome segregation protein |
| A01.119 | 633951 (+) | NCR | NA | - | NA, 274 bp upstream of MAG7520 (histone-like protein) |
| A01.275 | 447533 (+) | MAG3780 | 0.88 (+) | + | K03702 Excinuclease ABC subunit B |
| A02.068 | 600144 (+) | MAG5120 | 0.78 (+) | + | K00756 Pyrimidine-nucleoside phosphorylase |
| A03.187 | 581417 (+) | MAG4990 | 0.32 (+) | - | K01151 Deoxyribonuclease IV |
| A04.213 | 404407 (+) | MAG3410 | 0.56 (-) | + | No KEGG assignment (transposase) |
| A05.262 | 673860 (+) | MAG5810 | 0.05 (-) | + | No KEGG assignment (HP) |
| A06.084 | 289322 (+) | MAG2480 | 0.30 (-) | - | K09760 DNA recombination protein RmuC |
| A06.088 | 445824 (+) | MAG3780 | 0.03 (+) | + | K03702 Excinuclease ABC subunit B |
| A06.100 | 446738 (+) | MAG3780 | 0.49 (+) | + | K03702 Excinuclease ABC subunit B |
| A06.115 | 512566 (-) | MAG4380 | 0.32 (+) | - | K03529 Chromosome segregation protein |
| A06.302 | 011513 (+) | MAG0120 | 0.85 (-) | ++ | K07335 Basic membrane protein A (P48 lipoprotein) |
| A08.142 | 449880 (+) | MAG3790 | 0.74 (+) | + | K03701 Excinuclease ABC subunit A |
| A08.168 | 012048 (-) | MAG0120 | 0.53 (+) | ++ | K07335 Basic membrane protein A (P48 lipoprotein) |
| A09.087 | 011929 (-) | MAG0120 | 0.55 (+) | ++ | K07335 Basic membrane protein A (P48 lipoprotein) |
| A11.139 | 447945 (-) | MAG3790 | 0.06 (-) | + | K03701 Excinuclease ABC subunit A |
| A11.156 | 584228 (-) | MAG5020 | 0.08 (-) | - | K03703 Excinuclease ABC subunit C |
| A13.025 | 017476 (-) | MAG0160 | 0.50 (-) | + | K23536 Nucleoside transport system permease protein |

<sup>a</sup> Mutants were designated according to transformation and clone numbers (e.g., A01.088 designate clone 88 isolated from *M. agalactiae* transformation A01); <sup>b</sup> Insertion sites have been defined based on the sequence of PG2 (GenBank accession number: [CU179680.1](https://www.ncbi.nlm.nih.gov/nuccore/CU179680.1)). Orientation of the mTn is indicated in parenthesis using the puromycin-resistance maker as reference; <sup>c</sup> CDS accession number in the GenBank database; <sup>d</sup> Relative position and orientation of mTn in each interrupted CDS. HP, hypothetical protein; NA, Not applicable NCR; Non-coding region. <sup>e</sup> Level of resistance to ICEA invasion (see Fig. 1A, 72 h): -, none; +, intermediate; ++, high. <sup>f</sup> source: Kyoto Encyclopedia of Genes and Genomes (KEGG) functional orthology (KO) database (https://www.genome.jp/kegg/).

Resistance phenotypes were validated in cell culture mating assays using 24-well plates and controlled mycoplasma titers (Fig. 1A and Table 2). After 16 h of co-incubation, 9 mutants displayed resistance to ICE invasion. By 72 h, only 3 mutants (A06.302, A08.168 and A09.087) remained fully resistant. Remarkably, all these 3 mutants carried mTn insertions within MAG0120 (Fig. 1B), encoding the immunodominant surface lipoprotein P48.

**Figure 1.**
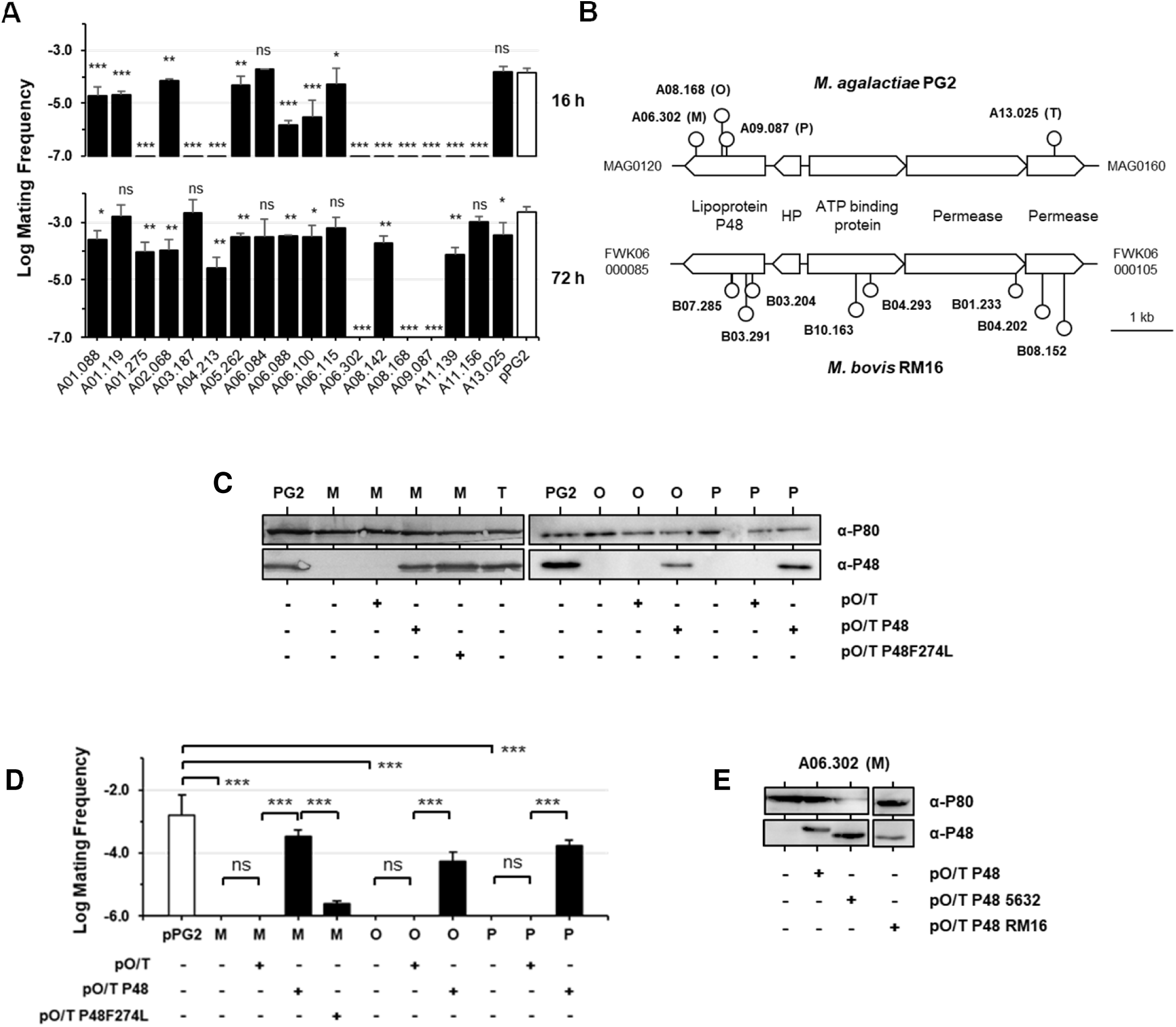
The ABC transporter-associated lipoprotein P48 is required for efficient ICE acquisition. (**A**) Resistance of *M. agalactiae* mutants to ICE transfer. Resistance was assessed by measuring the frequency of ICE transfer from PG2^T^[ICEA]^G^. The recipient pPG2^P^ (pPG2) was included as positive controls (Table 1). Mating frequency was calculated as the number of dual-resistant transconjugants per total CFUs. Data represent the mean of ≥3 independent experiments; error bars show standard deviations. The *p*-values were calculated with two-tailed t-tests relative to pPG2^P^. Significance: ns, *p* ≥ 0.05; *, *p* < 0.05; **, *p* < 0.01; ***, *p* < 0.001. (**B**) Transposon insertion map of the P48-associated ABC transporter locus in PG2 and RM16. The *P48* gene (MAG0120 in PG2; FWK06_000085 in RM16) is transcribed opposite to downstream ABC transporter genes encoding an ATP-binding protein and two permeases. Circles indicate insertion with corresponding mutant codes. (**C**) Western blot of P48 expression in the wild-type PG2 and P48 mutants A06.302 (M), A08.168 (O), A09.087 (P), and ABC transporter permease mutant A13.025 (T), with or without plasmids pO/T-P48, pO/T-P48F274L, or the empty vector pO/T. Lipoprotein P80 served as loading control. (**D**) Mating frequencies of P48 mutants and complemented strains Mating frequencies were calculated as the number of dual-resistant colonies per total CFUs. The pPG2^P^ pool (Table 1) was used as positive control. (**E**) Western blot of P48 expression in mutant A06.302 (M) complemented with pO/T-P48, pO/T-P48-5632, or pO/T-P48-RM16.

Thus, multiple chromosomal regions of the recipient cell can modulate ICEA transfer in *M. agalactiae*, but inactivation of the *p48* gene consistently conferred complete resistance, highlighting this locus as a critical determinant of ICEA acquisition.

### *M. bovis* genome-encoded determinants influencing ICEB acquisition

Given the close phylogenetic relationship between *M. agalactiae* and *M. bovis* (30), and the structural similarity of their ICEs (ICEA and ICEB, respectively) (18), we mapped *M. bovis* chromosomal regions involved in ICEB acquisition to identify recipient determinants of ICE transfer conserved in these two ruminant mycoplasmas.

We generated a transposon mutant library of 3,226 clones in the ICE-negative *M. bovis* strain RM16 and used J228[ICEB]^G^ as ICEB donor (Table 1). Unlike *M. agalactiae,* initial attempts to monitor ICEB transfer was unsuccessful with standard high-throughput conditions, likely reflecting the lower mating frequency of *M. bovis* (23). To overcome this, we modified the protocol by introducing a 24 h post-mating incubation in SP4 broth before plating.

This adapted screen identified 17 mutants with increased resistance to ICEB acquisition, each corresponding to a unique mTn insertion across 12 CDSs and one non-coding region (Table 3). Eight of the resistant mutants (B01.233, B03.204, B03.291, B04.202, B04.293, B07.285, B08.152, and B10.163) carried insertions within a conserved ABC transporter locus, including FWK06_000085 (Fig. 1B). This CDS encodes a lipoprotein showing 88.9% of overall similarity to *M. agalactiae* P48 (Fig. S1). The remaining insertions disrupted diverse genes not identified in the *M. agalactiae* screen, possibly reflecting differences in protocols or species-specific factors.

**Table 3.** Transposon insertion sites in selected *M. bovis* RM16 mutants.

| Mutant <sup>a</sup> | Genomic position <sup>b</sup> | CDS <sup>c</sup> | Relative position <sup>d</sup> | Resistance to ICEB <sup>e</sup> | KEGG orthology assignment <sup>f</sup> |
| --- | --- | --- | --- | --- | --- |
| B01.140 | 299265 (-) | NCR | NA | - | NA, 12 bp upstream of FWK06_001350 (HP) |
| B01.233 | 018503 (-) | FWK06_000100 | 96.9 (-) | ++ | K23535 nucleoside transport system permease protein |
| B02.157 | 022740 (-) | FWK06_000120 | 16.5 (+) | - | No KEGG assignment (lipoprotein, S41B peptidase family) |
| B03.204 | 013964 (+) | FWK06_000085 | 23.3 (-) | ++ | K07335 basic membrane protein A (P48 homolog) |
| B03.291 | 013868 (-) | FWK06_000085 | 30.1 (+) | + | K07335 basic membrane protein A (P48 homolog) |
| B04.202 | 018871 (-) | FWK06_000105 | 31.4 (-) | + | K23536 nucleoside transport system permease protein |
| B04.293 | 016059 (-) | FWK06_000095 | 67.3 (-) | ++ | K23537 nucleoside transport system ATP-binding protein |
| B07.285 | 013638 (+) | FWK06_000085 | 46.4 (-) | ++ | K07335 basic membrane protein A (P48 homolog) |
| B08.152 | 019277 (+) | FWK06_000105 | 73.0 (+) | ++ | K23536 nucleoside transport system permease protein |
| B09.044 | 585278 (+) | FWK06_002600 | 98.6 (+) | ++ | K01916 NAD+ synthase |
| B09.122 | 843729 (+) | FWK06_003700 | 94.6 (-) | + | K03168 DNA topoisomerase I |
| B09.142 | 119272 (+) | FWK06_000565 | 47.5 (-) | + | No assignment (membrane protein) |
| B09.240 | 064750 (-) | FWK06_000285 | 34.6 (+) | - | K03650 tRNA modification GTPase |
| B09.260 | 753661 (+) | FWK06_003280 | 47.4 (-) | + | K12573 ribonuclease R |
| B09.291 | 625698 (+) | FWK06_002805 | 93.0 (+) | - | K08483 phosphoenolpyruvate-protein phosphotransferase |
| B10.163 | 015792 (-) | FWK06_000095 | 50.7 (-) | + | K23537 nucleoside transport system ATP-binding protein |
| B11.173 | 490939 (-) | FWK06_002135 | 54.1 (+) | + | No KEGG assignment (membrane protein) |

<sup>a</sup> Mutants were designated according to transformation and clone numbers (See Table 2); <sup>b</sup>
Insertion sites have been defined based on the sequence of RM16 (GenBank accession

Resistance to ICEB acquisition was confirmed in cell culture mating assays. After 72 h, six mutants (B01.233, B03.204, B04.293, B07.285, B08.152 and B09.044) showed complete resistance to ICEB transfer (Fig. S2 and Table 3). Five carried insertions in the ABC transporter locus, including FWK06_000085 (P48 homolog), FWK06_000095 (ATP-binding protein), FWK06_000100 and FWK06_000105 (permeases) (Fig. 1B). The only insertion outside this locus disrupted a putative NAD⁺ synthetase (FWK06_002600) (Table 3). Remarkably, several ABC transporter mutants (Fig. 1B), including the P48 homolog mutant B03.291, retained partial susceptibility to ICEB transfer (Fig. S2 and Table 3).

Together, these genome-wide analyses in *M. agalactiae* and *M. bovis* demonstrate that the conserved lipoprotein P48 is a major determinant of ICE acquisition, as its inactivation provides near-complete protection against ICE invasion.

### Disruption of the P48 gene nearly abolishes ICEA acquisition without affecting transmission

The partial susceptibility of the *M. bovis* P48 mutant B03.291 to ICEB transfer prompted us to re-examine the resistance phenotype of the *M. agalactiae* P48 mutants under large-volume mating conditions. After 72 h of incubation in 75 cm² cell culture flasks, a small number of dual-resistant colonies were recovered with a mating frequency of 1.1 × 10^-7^. PCR confirmed that these colonies corresponded to P48 mutants that had acquired the G-tagged ICEA from donor strain PG2^T^[ICEA]^G^ (Fig. S3). Thus, despite the strong resistance of P48 mutants, ICEA acquisition was not completely abolished, indicating that alternative, low-frequency pathways may exist.

We next asked whether P48 is also required for ICEA transmission. For this, we tested the conjugative capacity of the ICEA-positive P48 mutant A06.302[ICEA] (Table 1), which was selected upon large-volume mating experiments. Remarkably, its mating frequency (9.9 ± 1.6 × 10⁻⁴) was comparable to that of the wild-type donor PG2^T^[ICEA]^G^. Thus, while disruption of the *P48* gene nearly abolishes ICEA acquisition, it has no effect on ICEA transmission.

### Disruption of the P48 gene induces transcriptional activation of a vestigial ICE

To investigate why P48 mutants resist ICEA invasion, we compared the whole transcriptomes of P48-negative and P48-positive cells. RNA-sequencing was performed on the wild-type PG2, the P48 mutant A06.302, and its complemented derivative CA06.302 (Table 1), all grown for 24 h in SP4 medium. Although growth in SP4 medium differs from cell-culture mating conditions, A06.302 retained its resistance phenotype (data not shown), and its growth curve showed only minor differences relative to the wild-type PG2 (Fig. S4).

Disruption of the *p48* gene led to significant transcriptional changes, with log₂ fold differences ranging from +1.21 to +1.87 and −1.06 to −2.33 among the top 50 differentially expressed genes (DEGs) (Table S1). As expected, *p48* gene expression itself was downregulated (Table S1), with almost no transcripts detected downstream of the insertion site (Fig. S5). Other genes downregulated relative to PG2 and CA06.302 were associated with redox metabolism and cellular homeostasis (glyceraldehyde-3-phosphate dehydrogenase, NADH oxidase, thioredoxin), as well as several uncharacterized products.

Remarkably, multiple genes within a vestigial ICEA (vICEA) region (MAG3860 to MAG4060) were upregulated in the mutant relative to PG2 (8 genes) and CA06.302 (3 genes) (Table S1). Additional upregulated genes included a cluster of four genes (MAG0710 to MAG0740), with two (MAG0710 and MAG0720), known to be essential for *M. agalactiae* proliferation in cell culture (31, 32).

These results show that disruption of the *p48* gene alters global gene expression, including activation of vICEA genes. Such transcriptional changes may create a cellular state unfavorable to a stable ICEA integration, potentially explaining the resistance phenotype. However, since ICEA-positive P48 mutants do not exhibit increased conjugative activity, their resistance is unlikely to result from ICEA instability.

### Lipoprotein P48 expression by recipient cells facilitates ICEA acquisition

The resistance of *M. agalactiae* P48 mutants to ICEA invasion prompted us confirm the role of P48 in ICEA acquisition. First, to rule out an influence of ICEA chromosomal location in the donor, we repeated mating assays using strain PG2^T^[ICEA]^G^-2 (Table 1), which differs from PG2^T^[ICEA]^G^ only in ICEA integration site. Mutant A06.302 remained resistant, indicating that the phenotype was independent of donor ICEA position.

Western blotting further confirmed the absence of P48 in three resistant mutants (A06.302, A08.168, A09.087) (Fig. 1C). Although anti-P48 antibodies detected truncated recombinant P48 fragments produced in *E. coli* (Fig. S6), no truncated protein was observed in the mutants, even when insertion sites were located in the central region or near the 3′ end of *p48* (Table 2). Attempts to express P48 truncated at Glu247 or Lys402, the codons disrupted by mTn insertions in A09.087 and A06.302, respectively, did not yield detectable protein products. These data indicate that no truncated forms of the P48 lipoprotein can be expressed in P48 mutants, reinforcing that functional P48 was absent from these mutants.

Complementation experiments provided definitive evidence of the role of P48 in ICEA acquisition. Transformation of P48 mutants with plasmid pO/T-P48 restored both P48 expression and susceptibility to ICEA transfer, whereas the empty vector had no effect (Fig. 1C-D). Finally, growth rates under mating conditions were unaffected, excluding fitness differences as an explanation for resistance (Fig. S4).

These results establish that expression of lipoprotein P48 in recipient cells is required for efficient ICEA acquisition, thereby confirming its role as a critical determinant of conjugation.

### Sequence variations in lipoprotein P48 influence ICEA acquisition

To test whether sequence variation in P48 affects ICEA transfer, we expressed P48 homologs from *M. agalactiae* 5632 (98.7% similarity) and *M. bovis* RM16 (88.9% similarity) in the P48 mutant A06.302 (Fig. S1). Western blotting confirmed successful expression of both homologs, although each displayed a slightly lower apparent molecular weight compared to PG2 P48 (Fig. 1E). This observation was unexpected since P48 in PG2 and 5632 have the same length, while P48 in PG2 and RM16 differ by 3 amino acids.

Mating experiments revealed that the 5632 P48-homolog restored ICEA acquisition to near wild-type levels (1.4 ± 0.4 × 10⁻³), whereas the RM16 homolog failed to complement the mutant, leaving cells resistant to ICEA transfer.

These findings demonstrate that sequence variations in P48 influence its ability to promote conjugation. Even small differences between homologs can markedly alter ICEA uptake, suggesting that the structural features required for conjugative transfer are highly specific.

### No detectable P48-CDS14 interaction in mating partner contact

ICEA-positive *M. agalactiae* cells constitutively express the ICEA-encoded surface lipoprotein CDS14, which is essential for conjugation (14). Since both CDS14 (donor-encoded) and P48 (recipient and donor-encoded) are surface-exposed lipoproteins required for ICEA transfer, we tested whether they interact during mating.

Reciprocal co-immunoprecipitation (co-IP) assays were performed with antibodies specific to P48 or CDS14 following co-incubation of the mating partners (PG2^T^[ICEA]^G^ donor and pPG2^P^ recipient; Table 1). Proteomic analysis of the anti-CDS14 eluate identified 234 proteins, with P48 ranking among the top 10 most abundant (Fig. 2A and Tables S2). By contrast, analysis of the anti-P48 eluate recovered 285 proteins but no detectable CDS14 peptides. Although both proteins were enriched in their respective pull-downs, no consistent evidence for a direct P48– CDS14 complex was obtained.

**Figure 2.**
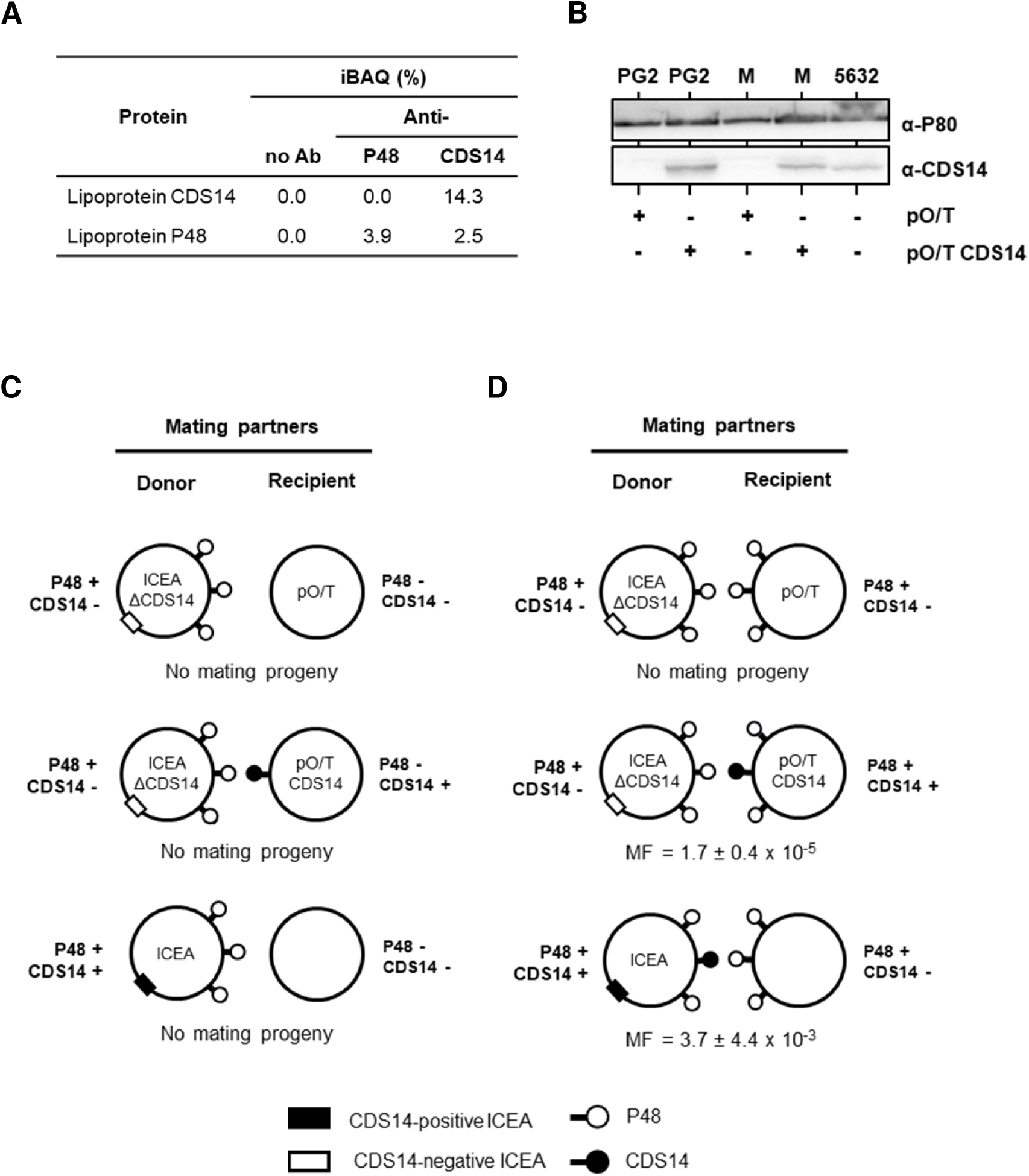
No detectable interaction between P48 and the ICEA-encoded surface lipoprotein CDS14. (**A**) Relative intensity-Based Absolute Quantification (iBAQ) values of CDS14 and P48 in co-IP fractions from mixed cultures of donor and recipient cells using anti-P48 or anti-CDS14 antibodies. (**B**) Western blot of CDS14 expression in PG2 and A06.302 (M) complemented with pO/T-CDS14 or empty vector. ICEA-positive strain 5632 served as CDS14-positive control. Lipoprotein P80 was used as loading control. (**C-D**) Schematic of mating experiments using a CDS14-negative ICEA donor and either (C) the P48 mutant A06.302 complemented with pO/T-CDS14 or the empty vector pO/T, or (D) the wild-type PG2 transformed with the same constructs. A CDS14-positive ICEA donor was included as control. MF, mating frequency (dual-resistant colonies per total CFUs).

In a previous study, we demonstrated that expression of CDS14 in the recipient can restore ICEA transfer from a CDS14-deficient donor (14). Thus, to further investigate the hypothesis of a P48-CDS14 interaction, we examined whether expressing CDS14 in a P48-deficient recipient (Fig. 2B) could rescue transfer from a CDS14-deficient but P48 positive donor (PG2^T^[ICEA *cds14::mTn*]^G^, Table 1). Mating experiments revealed no transfer when CDS14 was expressed in the P48 mutant A06.302 (Fig. 2C), while expression of CDS14 in wild-type recipients (Fig. 2B) restored ICEA transfer as expected (Fig. 2D).

Together, these data argue against a direct P48–CDS14 interaction in establishing donor– recipient contact, suggesting that P48 facilitates conjugation through alternative mechanisms.

### Lipoprotein P48 is a substrate-binding protein involved in nucleoside uptake

*In silico* analyses suggested that P48 functions as a substrate-binding protein for nucleoside uptake. This prediction is consistent with the presence of a Purine Nucleoside Receptor A (PnrA)-like domain (InterPro <u>IPR003760</u>), previously shown to mediate nucleoside uptake in other bacteria, including *Streptococcus pneumoniae* (33). It is also consistent with the identification downstream of *p48* (MAG0120) of several genes (MAG0140–MAG0160) encoding additional components of a putative ABC transporter (Fig. 1B). Structural predictions further supported this function. The AlphaFold model of P48 showed strong structural similarity to *S. pneumoniae* PnrA despite only 33.5% sequence identity (Fig. 3A and Fig. S7). The predicted P48 binding pocket conserved key residues involved in nucleoside recognition, including D78, F81, N82, D166, F274, G299, D323, and K341, which align with functionally equivalent residues in PnrA (Fig. 3A).

**Figure 3.**
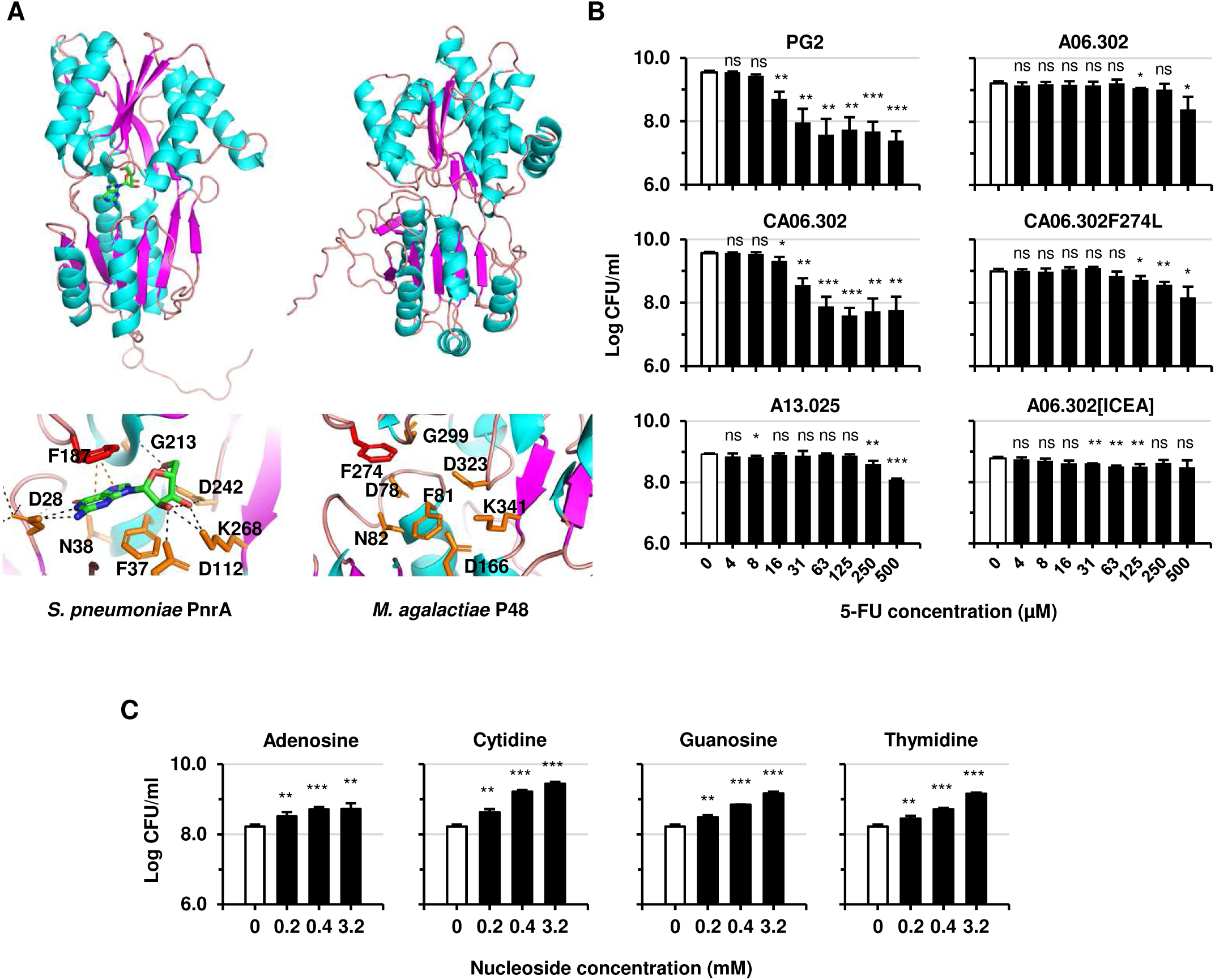
Lipoprotein P48 binds nucleosides and facilitates their transport via its associated ABC transporter. (**A**) Crystal structure of *S. pneumoniae* PnrA bound to guanosine (PDB: <u>6ya3</u>) and AlphaFold model of *M. agalactiae* P48 (<u>AF-F5HGV8</u>) with a mean pLDDT score of 90.19. Binding pocket residues in PnrA and corresponding residues in P48 (lower panels). Black and orange dashed lines represent polar and π-stacking interactions, respectively; the red stick highlights residue F274 mutated to leucine. (**B**) 5-FU toxicity assays with wild-type PG2, P48 mutant A06.302, complemented derivative CA06.302 (pO/T-P48) and CA06.302_ F274L (pO/T-P48F274L), the ABC transporter permease mutant A13.025, and the ICEA-positive P48 mutant A06.302[ICEA]. Titers were measured after 24 h in SP4 with increasing 5-FU concentrations. (**C**) Nucleoside competition assay. PG2 titers after 24 h in SP4 with 0.3 mM 5-FU alone (open bars) or combined with adenosine, cytidine, guanosine, or thymidine. Bars show are means of ≥3 independent experiments. Error bars represent standard deviations. Significance: ns, *p* ≥ 0.05; *, *p* < 0.05; **, *p* < 0.01; ***, *p* < 0.001.

Functional assays using the toxic nucleoside analogue 5-fluorouridine (5-FU) confirmed that P48 functions as a substrate-binding protein involved in nucleoside uptake. Wild-type PG2 cells were highly sensitive to 5-FU, whereas P48 mutant A06.302 was resistant. Complementation with plasmid pO/T-P48 restored 5-FU sensitivity, thereby confirming the role of P48 in nucleoside uptake (Fig. 3B). Competition assays with natural nucleosides further showed that adenosine, guanosine, cytidine, and thymidine reduced 5-FU toxicity, indicating broad substrate promiscuity (Fig. 3C).

These results demonstrate that P48 is a nucleoside-binding lipoprotein and a substrate-binding component of an ABC transporter. The broad nucleoside recognition capacity of P48 underscores its key role in mediating *M. agalactiae* interactions with the extracellular environment to facilitate nutrient acquisition.

### The nucleoside-binding activity of lipoprotein P48 is required for efficient ICE acquisition

To test whether nucleoside-binding by P48 is necessary for ICEA acquisition, we engineered a point mutation in the predicted binding pocket. Phenylalanine 274, a residue expected to mediate π-stacking interactions with nucleosides, was substituted with leucine (F274L) to disrupt binding while maintaining hydrophobicity and side-chain size (Fig. 3A).

Expression of the mutant protein was confirmed by Western blotting (Fig. 1C). Functional assays showed that cells expressing P48-F274L were resistant to 5-FU, indicating impaired nucleoside binding, whereas wild-type P48 restored sensitivity (Fig. 3B).

Mating assays revealed that complementation of the P48 mutant A06.302 with P48-F274L failed to restore ICEA acquisition to wild-type levels (Fig. 1D).

These findings demonstrate that the nucleoside-binding activity of P48 is essential for efficient ICEA acquisition suggesting a functional link between nucleoside recognition and ICEA transfer in *M. agalactiae*.

### ICEA acquisition relies on P48 nucleoside-binding activity but not on nucleoside uptake

Lipoprotein P48 is predicted to function as the substrate-binding component of an ABC transporter that also includes an ATP-binding protein (MAG0140) and two permeases (MAG0150, MAG0160) (Fig. 1B). To test whether nucleoside transport itself was required for ICEA transfer, we analyzed mutant A13.025, in which MAG0160 was disrupted (Fig. 1B and Table 2).

As expected for loss of a transporter permease, mutant A13.025 was resistant to 5-FU toxicity, reaching tolerance levels comparable to the P48 mutant A06.302 (Fig. 3B). Western blotting confirmed that P48 expression was intact in A13.025, ruling out secondary effects on protein stability (Fig. 1C).

Remarkably, despite its inability to import nucleosides, mutant A13.025 remained permissive to ICEA acquisition in mating assays (Fig. 1A). This indicates that nucleoside transport in PG2 is dispensable for conjugative transfer, and that ICEA acquisition relies specifically on the binding activity of P48 rather than its transport function.

It is noteworthy that *M. bovis* mutants with disrupted ABC transporter genes all displayed partial or complete resistance to ICEB transfer (Fig. S2 and Table 3). This suggests that the mechanisms underlying ICEB dissemination in *M. bovis* may differ from those governing ICEA transfer in *M. agalactiae*, despite the close phylogenetic relationship between these two ruminant mycoplasma species.

## DISCUSSION

Integrative and conjugative elements (ICEs) are self-transmissible and have long been studied from a donor-centric perspective, with emphasis on ICE-encoded functions and the conjugative machinery. Recipient cells, in contrast, were largely regarded as passive partners, their role limited to mounting defense systems such as restriction–modification, CRISPR-Cas, or the SOS response that restrict horizontal gene flow (1). Beyond these classical post-entry barriers, only a few recipient-encoded factors have been shown to modulate conjugative transfer (21, 28, 29).

Our study challenges this paradigm by demonstrating that ICE transfer in mycoplasmas is tightly dependent on a recipient-encoded lipoprotein, and specifically on its nucleoside-binding activity. This finding highlights the active contribution of recipient functions to ICE dissemination and suggests that ICE acquisition may be selectively regulated at the recipient cell surface.

### Recipient determinants of ICE acquisition

While P48 emerged as the primary determinant of ICE acquisition in *M. agalactiae*, our genome-wide screen also uncovered additional putatively contributing loci, including genes involved in DNA repair (e.g., *uvrABC*), nucleoside metabolism, and general DNA processing. While these disruptions only partially affected ICE acquisition, they suggest that a broader network of recipient pathways can modulate conjugation efficiency. Some of these genes, such as *uvrA*, also influence mycoplasma growth in cell culture (31, 32), complicating interpretation of their role in ICE transfer.

The implication of the recipient lipoprotein P48 in ICE acquisition was further confirmed in *M. bovis*. However, notable differences emerged in the recipient-encoded factors influencing ICE uptake. For instance, genes linked with DNA-related functions, such as *uvrABC*, were not identified in the *M. bovis* mutant screen. Another striking difference between the two species concerned the ABC transporter associated with lipoprotein P48. While this ABC transporter was found contributing to ICEB acquisition in *M. bovis*, it appeared to be largely dispensable in *M. agalactiae*. This difference may reflect species-specific variations or the methodological adjustments required by differences in mating efficiency. Regardless, the convergent identification of P48 in both species firmly establishes it as a key host factor for ICE dissemination.

### P48 as a nucleoside-binding lipoprotein

Because mycoplasmas lack a cell wall, lipoproteins such as P48 are directly exposed to the extracellular environment and are highly antigenic (34–36). Our work strongly suggests that P48 functions as a substrate-binding protein of an ABC transporter and mediates broad nucleoside recognition. This finding confirms earlier predictions for its *M. bovis* homolog (37). Remarkably, however, the role of P48 in conjugation was not linked to its transport function but specifically to its nucleoside-binding activity. This distinction was evident from two complementary observations. First, a point mutation disrupting the nucleoside-binding pocket (F274L) abolished ICEA acquisition despite normal expression of P48. Second, disruption of the cognate permease prevented nucleoside uptake but did not impair ICEA transfer. Together, these results establish that ICE acquisition depends on binding rather than transport, suggesting that P48 plays a signaling or structural role during conjugation.

P48 is highly conserved across members of the *Hominis* phylogenetic group, with the notable exception of *M. hominis*, a species that nonetheless carries ICEs (22, 38, 39). This observation implies that *M. hominis* relies on alternative mechanisms for conjugative uptake, potentially involving nucleoside-binding lipoproteins lacking a PnrA-like domain or other substrate-binding proteins associated with the candidate nucleoside ABC transporter protein BcrA (40). Whether such lipoproteins perform functions analogous to P48 in facilitating ICE transfer remains an open question.

### One protein, two possible mechanisms

The surface localization and nucleoside-binding activity of P48 suggest two nonexclusive ways it may facilitate ICEA acquisition.

First, P48 may act as a structural partner of the conjugative machinery. Because it is surface-exposed and constitutively expressed, P48 could contribute to stabilizing donor–recipient contact during transfer. A natural candidate for interaction is the ICEA-encoded lipoprotein CDS14, also present at the donor surface and essential for transfer (14). However, neither co-IP nor functional complementation supported a direct P48–CDS14 interaction. Interestingly, a cooperative exchange of DNA, involving both donor and recipient cells, has been recently discovered in *Thermus thermophilus* (41–45). This process couples the ICE-encoded DNA donation machinery of the donor with the natural competence system of the recipient. Although no natural competence system has been described in mycoplasmas, it remains possible that P48 could mediate analogous interactions, potentially accommodating nucleic acids in addition to nucleosides.

Second, P48 may function as a sensory protein. Its similarity to Med, a PnrA-like protein in *Bacillus subtilis* that promotes competence by activating the transcriptional regulator ComK (46–50), raises the possibility that nucleoside binding by P48 transduces a signal that promotes ICE acquisition. Yet, mycoplasmas lack classical competence pathways, making this scenario less likely. It should be noted that activation of the sigma factor σ^20^ in *M. genitalium* has been associated with an unusual form of HGT (26). However, this DNA transfer mechanism operates independently of any known mobile genetic elements. Alternatively, P48 signaling could influence ICEA stability post-integration. Indeed, disruption of p48 was associated with transcriptional activation of a vestigial ICEA. In this scenario, P48 would not be involved in early steps of ICEA acquisition, and the resistance of P48 mutants could instead result from instability of the chromosomal ICEA after its integration. Still, because ICEA-positive P48 mutants were viable and not hyper-conjugative, this effect is unlikely to fully explain the resistance phenotype.

Overall, both hypotheses remain plausible: P48 may act either as a structural facilitator of donor–recipient contact or as a sensor modulating cellular states permissive for ICE acquisition

### Nucleoside metabolism and ICE dissemination

Mycoplasmas are metabolically constrained and rely heavily on their hosts for essential nutrients (9, 10). In this context, nucleosides and nucleotides are increasingly recognized as central to their physiology. For example, their metabolism is increasingly recognized as a critical driver of *M. bovis* survival and virulence (51–54). Our previous work also showed that elevated deoxynucleotide concentrations inhibit ICEA transfer in *M. agalactiae* (23), suggesting that nucleotide availability may modulates conjugation.

The discovery that P48-mediated nucleoside binding is essential for ICE acquisition adds a new layer to this relationship. Beyond serving as metabolic substrates, nucleosides may act as extracellular cues that couple environmental conditions to HGT. In this view, mycoplasmas could exploit host-derived metabolites to fine-tune their conjugation activity, linking nutrient sensing directly to genome evolution.

### A target for the control of horizontal gene transfer in mycoplasmas

Synthetic biology has positioned live mycoplasmas as promising platforms for therapeutic applications (55–57). However, the safe deployment of engineered strains depends on maintaining genetic stability and limiting HGT. Because P48 is both surface-exposed and essential for conjugation, it represents a potential target to block ICE dissemination. This vulnerability at the mycoplasma cell surface, may prove valuable for controlling the spread of mobile genetic elements in pathogenic mycoplasmas and for safeguarding synthetic strains intended for biotechnological or medical use.

## Conclusion

This study reshapes our understanding of bacterial conjugation by demonstrating that recipient cells can actively determine the outcome of ICE transfer. Using *M. agalactiae* as a model, we demonstrate that ICE acquisition depends on the nucleoside-binding activity of lipoprotein P48, uncovering a critical surface function for conjugation. This challenges the donor-centric view of conjugation, positions nucleosides as metabolic signals for HGT, and open avenues to control ICE spread in pathogenic mycoplasmas and protect engineered strains.

## MATERIALS AND METHODS

### Mycoplasmas, cell lines and culture conditions

Antibiotic-tagged *M. agalactiae* and *M. bovis* strains used in this study are described in Table 1. Mycoplasmas were grown at 37°C in SP4 medium (58) supplemented with 100 µg/ml ampicillin (Sigma-Aldrich). When needed, gentamicin (50 µg/ml; Gibco), puromycin (10 µg/ml; Thermo Fisher), or tetracycline (2 µg/ml; Sigma-Aldrich) were added to the medium, alone or in combination. Stock cultures were stored at −80°C. Since mycoplasma growth cannot be monitored by optical density, titers were determined based on colony counts (CFU/ml) on solid SP4 media after 4-7 days of incubation at 37°C by using a binocular stereoscopic microscope. The detection limit for mycoplasma titration was 100 CFU/ml, except for the detection of mating progeny, which was 10 CFU/ml. T-antigen immortalized goat milk epithelial cells (TiGMEC) were kindly provided by C. Leroux (UMR754 IVPC, Lyon). Cells were grown in Dulbecco’s modified Eagle’s medium (DMEM, high glucose, sodium pyruvate, and GlutaMAX-I; Gibco) supplemented with non-essential amino acids (NEAA, Gibco) and 15% heat-inactivated fetal bovine serum (FBS, Gibco). The mycoplasma-free status of the cell line was routinely tested by genus-specific PCR (59).

### Random transposon mutagenesis and mapping of insertion sites

Random transposon mutant libraries were generated as previously described (32). Briefly, a puromycin-resistance tag was randomly inserted into the mycoplasma genome by transformation with plasmid pMT85Pur2 (23), a modified version of Tn*4001* (mTn) carrying the *pac* gene (60). For each library, up to 3000 individual colonies were collected from independent transformations and grown in 1 ml of selective SP4 medium. Cultures of individual mutants were distributed in 96-well plates and stored at −80°C. Transposon insertion sites were mapped by sequencing the junction between the chromosomal DNA and the 3’ end of the mTn. Genomic DNA from mutant cultures was amplified by single-primer PCR (61) using an mTn-specific primer SG6_2 (Table S3). PCR products were sequenced (Eurofins Genomics) with nested primers M13-24b_Rev, M13-30bases_REV or SG5 (Table S3).

### High-throughput mating assays

High-throughput mating experiments were conducted in 96-well plates (Falcon) under cell culture conditions. TiGMEC cells were seeded at a density of 4 × 10^4^ cells/cm^2^ and inoculated with each mating partner using the 96-pin replicator (Boekel Scientific). After 72 h incubation at 37°C in a humidified 5% CO_2_ atmosphere, the mating progeny, consisting of the P-tagged mutants having acquired the G-tagged ICE from the donor strains, were seeded on SP4 solid media supplemented with gentamicin (50 µg/ml; Gibco) and puromycin (10 µg/ml; Thermo Fisher). For *M. bovis*, the protocol included an additional 24 h post-mating incubation step. At the end of the initial 72 h incubation, an equal volume of fresh SP4 medium was added to each well, and plates were incubated for another 24 h before plating on selective media.

### Cell culture mating assays

Cell culture mating experiments were conducted in 24-well plates (Falcon) as previously described (23). TiGMEC cells were seeded at a density of 4 × 10^4^ cells/cm^2^ and inoculated with 10^6^ CFU/ml of each partner, followed by incubation for 16 or 72 h at 37°C in a humidified 5% CO_2_ atmosphere. After three freeze-thaw cycles, mycoplasma titers were determined by CFU titration on selective media, or without any selective pressure for total cell counts. Mating frequency was expressed as the number of dually resistant colonies divided by the total CFUs.

### DNA constructs for protein expression in mycoplasmas

Plasmid p20-1miniO/T (further designated pO/T) was used as an expression vector for complementation studies (32). To generate pO/T-P48, the *M. agalactiae* lipoprotein P48 gene (MAG0120) was cloned downstream of the *P40* promoter (32). The promoter region was amplified using primers p40RF-CC and P40/P48 N-ter (Table S3) to produce a 200 bp fragment overlapping *MAG0120* at the ATG start codon. *MAG0120* was then amplified using this overlapping fragment and primer P48 C-ter (Table S3). A similar strategy was used to construct pO/T-P48-5632, pO/T-P48-RM16, and pO/T-CDS14, which express the P48 homologs from *M. agalactiae* 5632 (MAGa0140) and *M. bovis* RM16 (FWK06_000085), and the ICEA CDS14 protein (MAGa3160), respectively. Plasmid pO/T-P48F274L was generated by introducing a single nucleotide substitution (T822A) in MAG0120, resulting in a Phe274→Leu substitution in P48. This substitution was introduced by PCR-based site-directed mutagenesis using mismatched overlapping oligonucleotide primers T822AF and T822AR (Table S3). Using the same approach, plasmids pO/T-P48A1204T and pO/T-P48C739T were constructed by introducing A1204T or C739T substitutions, respectively, converting Lys402 (AAA) and Gln247 (CAA) codons into stop codons (TAA). Mismatched overlapping oligonucleotide primers P48A1204T-F/-R and P48C739T-F/-R used for these constructs are listed in Table S3. PCR amplifications were performed using a high-fidelity DNA polymerase (Phusion; New England Biolabs), and all constructs were verified by Sanger sequencing (Eurofins Genomics). Mycoplasma transformation with plasmid DNA was carried out as described (32).

### Western blotting

Mycoplasma cultures were centrifuged at 10,000 x *g* for 20 min at 4°C. Pellets were washed and resuspended in DPBS before resuspension in Laemmli sample buffer. Western blotting was performed as described (62), with Samples were normalized to 2 × 10^8^ CFU per well. Anti-P48 serum was generated by rabbit immunization (Eurogentec) using recombinant P48 produced with the pMAL protein fusion and purification system (New England Biolabs). A sheep serum raised against *M. agalactiae* P80 surface antigen was used as a control (62). Blots were developed with HRP-conjugated swine anti-rabbit or rabbit anti-sheep IgG (DAKO) and visualized using Western Bright ECL (Advansta).

### 5-fluorouridine inhibition and nucleoside competition assays

Mycoplasmas were grown in SP4 medium containing with increasing concentrations of 5-fluorouridine (5-FU; Thermo Fisher). For competition assays, 10^7^ CFU were seeded in SP4 medium supplemented with 0.3 mM of 5-FU, either alone or in combination with increasing concentration of individual nucleosides (guanosine, adenosine, thymidine and cytidine; Thermo Fisher). After 24 h of incubation, growth was determined by CFU titration.

### Co-immunoprecipitation (Co-IP) assays

Mating partners were grown individually for 48 h in 300 ml of SP4 medium, pooled, and further incubated overnight at 4°C. Cells were harvested by centrifugation (12,000 x *g*, 20 min, 4°C), washed three times in DPBS (Invitrogen), and lysed in ice-cold lysis buffer (DPBS, 1% Triton X-114). Protein A-Sepharose CL-4B beads (0.5 g; Cytiva) were functionalized with anti-P48 or anti-CDS14 rabbit sera (14). Co-IP experiments were performed using. Beads were incubated with sera for 3 h at room temperature, washed with DPBS, and equilibrated in lysis buffer. Protein A–Sepharose beads without antibody were used as a negative control. Cell lysates were loaded onto the columns, and unbound proteins were removed by extensive washing with ice-cold lysis buffer. Bound proteins were eluted with ice-cold acidic buffer (0.1 M glycine HCl, 154 mM NaCl, 1% Triton X-114, pH 2.7), neutralized, and subjected to Triton X-114 phase separation and precipitation as described (62). Protein pellets were resuspended in Laemmli sample buffer and analyzed by SDS–PAGE and mass spectrometry.

### Proteomic analysis, data-Independent acquisition, and data processing

Proteins in 5% SDS were reduced and alkylated (100 mM TCEP, 400 mM chloroacetamide, 5 min, 95 °C), acidified (2.5% phosphoric acid), and diluted 7× in S-Trap buffer (90% methanol, 100 mM triethylammonium bicarbonate, pH 7.1). Samples were loaded onto S-Trap Micro spin columns (Protifi), washed three times with S-Trap buffer, and digested overnight at 37 °C with 4 µg trypsin in 50 mM ammonium bicarbonate. Peptides were sequentially eluted with (i) 50 mM ammonium bicarbonate, (ii) 0.2% formic acid, and (iii) acetonitrile/formic acid (50/0.2), dried, and resuspended in 0.2% formic acid. Peptides (1 µl) were separated on a heated PEPMAP C18 column (75 µm × 500 mm, 45 °C) using a 95 min gradient (0–25% B: 58 min; 25–40%: 20 min; 40–90%: 2 min; 90%: 5 min). Data were acquired on an Orbitrap Exploris 480 in DIA mode (MS1: 60,000 resolution, AGC 300%, 100 ms IT, 400–1008 m/z; MS2: 15,000 resolution, 2×75 staggered windows of 8 m/z, NCE 30). Raw files were processed with DIA-NN v1.8.1 (library-free search) against *M. agalactiae* PG2 and ICEA databases (CU179680.1, CT030003.1) and imported into Proline v2.1.2. Carbamidomethylation (C) was fixed; methionine oxidation was variable. Trypsin/P specificity allowed up to two missed cleavages. Peptide length was set to 7–30 aa, precursor m/z 400–1000, fragment ions 140–2000 m/z, with precursor FDR at 1.

### RNA isolation and RNA-sequencing

Mycoplasmas grown for 24 h in SP4 medium were harvested by centrifugation (12,000 × *g,* 5 min, 4°C). Total RNA was extracted from cell pellets (3 × 10^9^ CFUs) using TRIzol reagent (Invitrogen) and purified with RNAClean XP beads (Beckman Coulter). RNA concentration was measured spectrophotometrically (CLARIOstar Plus, BMG LABTECH), and RNA integrity was assessed using the Agilent RNA ScreenTape kit on a 4200 TapeStation (Agilent Technologies). cDNA library preparation and sequencing were performed at IGATech (Italy) using the Zymo-Seq RiboFree Total RNA library prep kit (Zymo). Libraries were quality checked with a Qubit 2.0 Fluorometer (Invitrogen) and Agilent Bioanalyzer DNA assay (or Caliper, PerkinElmer), then sequenced (paired-end, 150 bp) on a NovaSeq 6000 platform (Illumina). Raw reads were quality filtered using Trim Galore, and paired-end reads were aligned to the *M. agalactiae* PG2 genome (<u>CU179680.1</u>) using the Salmon v. 1.4. Count tables were imported using *tximport* function (tximport v.1.28) and analysis using the DESeq2 v. 1.40.2. *P*-values were adjusted for multiple testing using the Benjamini-Hochberg procedure (false discovery rate). Exploratory analysis of the expressed genes matrix was performed with principal component analysis (PCA) and multidimensional scaling (MDS) after the regularized-logarithm transformation (edgeR) or variance-stabilizing transformation in DESeq2.

## Data Availability Statement

The mass spectrometry proteomic data have been deposited to the ProteomeXchange Consortium via the PRIDE partner repository with the dataset identifier PXD068569. The RNA-seq data generated for this study have been deposited in the European Nucleotide Archive (ENA) under the project accession number PRJEB97083. Individual sample accessions are as follows: ERS26772405 to ERS26772407 (PG2 A06.302_1 to 3); ERS26772408 to ERS26772410 (PG2 CA06.302_1 to 3); ERS26772411 to ERS26772413 (PG2_1 to 3). All associated FASTQ files have been uploaded and are publicly accessible.

## Acknowledgments

We thank R. Herrmann for providing plasmid pMT85 and C. Leroux for TiGMEC cells. We also thank IGA Technology Services (www.igatechnology.com; Udine, Italy) for providing sequencing support and infrastructure for the analysis of our RNA samples and H. Lebrette for helpful discussions.

## Author Contributions

All authors read and approved the version submitted for publication.

**A. M. Derriche:** Conceptualization, Data curation, Formal analysis, Investigation, Validation, Visualization, Writing - original draft.

**LX. Nouvel:** Conceptualization, Data curation, Formal analysis, Methodology, Supervision, Validation, Writing - review & editing.

**Calvin Fauvet:** Investigation, Methodology, Validation.

**Nuria Mach:** Data curation, Formal analysis.

**E. Simon:** Investigation, Methodology, Validation.

**G. Pot:** Investigation, Methodology, Validation.

**H. Robert:** Investigation, Methodology, Validation.

**A. Stella**: Investigation, Methodology, Validation.

**B. de la Fe:** Funding acquisition, Resources.

**R. Maillard:** Funding acquisition, Resources.

**Sergi Torres-Puig:** Conceptualization, Writing - review & editing.

**Y. Arfi:** Funding acquisition, Writing - review & editing.

**C. Citti:** Conceptualization, Funding acquisition, Project administration, Writing - review & editing.

**E. Baranowski:** Conceptualization, Data curation, Funding acquisition, Methodology, Project administration, Resources, Supervision, Validation, Visualization, Writing - original draft, Writing - review & editing.

## SUPPLEMENTARY TABLES

**Table S1.** Top 50 DEGs in the P48 mutant A06.302.

| A06.302 vs PG2<sup>a</sup> |  |  | A06.302 vs CA06.302<sup>a</sup> |  |  |
| --- | --- | --- | --- | --- | --- |
| Gene<sup>b</sup> | FC<sup>c</sup><br>(Log<sub>2</sub>) | UniProtKB/TrEMBL<sup>d</sup> | Gene<sup>b</sup> | FC<sup>c</sup><br>(Log<sub>2</sub>) | UniProtKB/TrEMBL<sup>d</sup> |
| *Upregulated genes* |  |  |  |  |  |
| **MAG1560** | **1.87** | **Uncharacterized lipoprotein** | MAG2188 | 2.38 | No entry (tRNA-Phe) |
| **MAG4040** | **1.78** | **VirD4/TraG family (vICEA)** | MAG7100 | 2.00 | Variable surface lipoprotein |
| MAG0300 | 1.74 | Uncharacterized protein | MAG0657 | 1.74 | No entry (tRNA-Arg) |
| MAG1120 | 1.73 | No entry (pseudogene) | MAG0700 | 1.35 | Serine hydroxymethyltransferase |
| **MAG4060** | **1.73** | **Uncharacterized protein (vICEA)** | MAG0660 | 1.30 | ATPase, AAA family |
| MAG0730 | 1.69 | Nitrogen fixation protein NifU | **MAG1560** | **1.29** | **Uncharacterized lipoprotein** |
| **MAG4030** | **1.67** | **C5 Mtase (vICEA)** | MAG2190 | 1.28 | Cytidine triphosphate synthase |
| MAG0740 | 1.65 | DNA damage repair protein MucB | **MAG4030** | **1.24** | **C5 Mtase (vICEA)** |
| MAG2910 | 1.64 | Uncharacterized protein | **MAG0710** | **1.22** | **Proline iminopeptidase** |
| MAG4050 | 1.61 | Uncharacterized protein (vICEA) | MAG0670 | 1.19 | Phenylalanine--tRNA ligase PheS |
| MAG1140 | 1.60 | Uncharacterized protein | **MAG4040** | **1.12** | **VirD4/TraG family (vICEA)** |
| MAG4070 | 1.60 | Uncharacterized protein | MAG0690 | 1.11 | Phenylalanine--tRNA ligase PheT |
| **MAG0720** | **1.60** | **Nitrogen fixation protein NifS** | **MAG0310** | **1.10** | **Uncharacterized protein** |
| **MAG6150** | **1.50** | **No entry (pseudogene)** | **MAG4060** | **1.07** | **Uncharacterized protein (vICEA)** |
| MAG2900 | 1.49 | Uncharacterized protein | MAG6780 | 1.07 | tRNA methyltransferase |
| MAG6160 | 1.47 | No entry (pseudogene) | MAG0680 | 1.02 | Uracil-DNA glycosylase |
| MAG4010 | 1.45 | IrrE N-term-like domain-containing protein (vICEA) | MAG6010 | 0.98 | Uncharacterized protein |
| MAG3990 | 1.44 | ssDNA-binding protein (vICEA) | **MAG0720** | **0.96** | **Nitrogen fixation protein NifS** |
| **MAG0310** | **1.39** | **Uncharacterized protein** | MAG5030 | 0.94 | Lipoprotein P80 |
| MAG0330 | 1.33 | Uncharacterized protein | MAG6660 | 0.90 | Dephospho-CoA kinase |
| MAG3900 | 1.29 | Uncharacterized protein (vICEA) | MAG7110 | 0.90 | Integrase recombinase Xer |
| **MAG0710** | **1.27** | **Proline iminopeptidase** | MAG0400 | 0.86 | CysteinyI tRNA synthetase |
| MAG2920 | 1.23 | Transmembrane protein | MAG2240 | 0.83 | Holliday junction branch migration complex SU RuvB |
| MAG6880 | 1.23 | Uncharacterized protein | **MAG6150** | **0.83** | **No entry (pseudogene)** |
| MAG3980 | 1.21 | Uncharacterized protein (vICEA) | MAG2560 | 0.83 | Pseudogene |
| *Downregulated genes* |  |  |  |  |  |
| **MAG0550** | **-2.33** | **Glyceraldehyde-3-phosphate dehydrogenase** | MAG3190 | -1.94 | Enolase |
| **MAG2630** | **-1.78** | **NADH oxidase** | MAG2010 | -1.90 | Large ribosomal SU protein bL33B |
| MAG1460 | -1.77 | Chaperone protein DnaK | **MAG0550** | **-1.59** | **Glyceraldehyde-3-phosphate dehydrogenase** |
| MAG3510 | -1.74 | ATP synthase subunit c | MAG5880 | -1.23 | Transmembrane protein |
| **MAG7380** | **-1.63** | **Thioredoxin** | **MAG7380** | **-1.20** | **Thioredoxin** |
| MAG2740 | -1.56 | Alcohol dehydrogenase | MAG4180 | -1.07 | Uridine phosphorylase |
| MAG3500 | -1.52 | ATP synthase A chain | MAG4910 | -1.05 | Lipoyl-binding domain-containing protein |
| MAG5110 | -1.47 | Uridine phosphorylase | MAG4560 | -0.98 | Small ribosomal SU protein uS9 |
| **MAG5870** | **-1.38** | **Uncharacterized protein** | **MAG5870** | **-0.90** | **Uncharacterized protein** |
| MAG7250 | -1.37 | Cytidylate kinase | MAG0020 | -0.88 | DNA polymerase III, beta chain |
| MAG0775 | -1.21 | HTH HARE-type domain- containing protein | MAG4680 | -0.88 | Acyl carrier protein phosphodiesterase |
| MAG0100 | -1.17 | Luciferase-like domain-containing protein | MAG2350 | -0.88 | Variable surface lipoprotein |
| MAG0160 | -1.15 | Sugar ABC transporter permease | **MAG0120** | **-0.87** | **Lipoprotein P48** |
| MAG3960 | -1.15 | Uncharacterized protein | MAG6450 | -0.87 | Uncharacterized protein |
| MAG3520 | -1.14 | ATP synthase subunit b | **MAG2630** | **-0.85** | **NADH oxidase** |
| MAG4900 | -1.12 | L lactate dehydrogenase | MAG0010 | -0.83 | Chromosomal replication initiator protein DnaA |
| MAG4280 | -1.12 | Alcohol dehydrogenase | MAG6180 | -0.80 | Large ribosomal SU protein bL12 |
| **MAG2220** | **-1.12** | **Uncharacterized lipoprotein** | MAG4530 | -0.77 | Ion trans domain protein |
| MAG6370 | -1.11 | Pentitol phosphotransferase enzyme II, B component | MAG2320 | -0.77 | Glycerol ABC transporter permease gtsB |
| **MAG0120** | **-1.10** | **Lipoprotein P48** | **MAG2220** | **-0.76** | **Uncharacterized lipoprotein** |
| MAG4690 | -1.10 | Phosphocarrier protein HPr | MAG5140 | -0.75 | Triosephosphate isomerase |
| MAG5120 | -1.06 | Thymidine phosphorylase | MAG2330 | -0.75 | Glycerol ABC transporter permease gtcC |
| MAG3540 | -1.06 | ATP synthase subunit alpha | MAG4190 | -0.73 | Uncharacterized protein |
| MAG3460 | -1.06 | Uncharacterized protein | MAG4970 | -0.71 | Hexosephosphate transport protein |
| MAG3050 | -1.06 | Uncharacterized protein | MAG6400 | -0.70 | Uncharacterized protein |

<sup>a</sup> RNA sequencing was performed on three biological replicates; DEG with a *p*-value < 0.01 and adjusted *p*-value < 0.01; CA06.302 designates the complemented derivative of the P48 mutant A06.302; entries identified in both comparative analysis are in bold letters; vICEA: genes associated with the vestigial ICEA identified in the PG2 genome (18); <sup>b</sup> CDS identifier from the PG2 genome in the GenBank database ([CU179680.1](https://www.ncbi.nlm.nih.gov/nuccore/CU179680.1)); <sup>c</sup> Log<sub>2</sub> Fold Change (FC) in RNA levels; <sup>d</sup> Protein names according to UniProtKB/TrEMBL entries in the UniProt Knowledgebase (UniProtKB; https://www.uniprot.org/).

**Table S2.** Top 10 proteins immunoprecipitated with anti-P48 and anti-CDS14 antibodies ^a^.

| Proteins <sup>b</sup> | CDS <sup>c</sup> | IBAQ (%) <sup>d</sup> |
| --- | --- | --- |
| *Proteins co-IP with P48 antibodies* |  |  |
| Variable surface lipoprotein Y | MAG7080 | 12.86 |
| Variable surface lipoprotein | MAG3600 | 7.14 |
| Variable surface lipoprotein X | MAG7070 | 5.90 |
| Lipoprotein P48 | MAG0120 | 3.92 |
| LemA family protein | MAG1220 | 3.67 |
| Small ribosomal subunit protein uS3 | MAG5400 | 3.12 |
| NADH oxidase | MAG2630 | 2.78 |
| ATP-dependent zinc metalloprotease | MAG7450 | 2.39 |
| Lipoprotein P80 | MAG5030 | 2.26 |
| Hexosephosphate transport protein | MAG4970 | 2.15 |
| *Protein co-IP with CDS14 antibodies* |  |  |
| Lipoprotein CDS14 | cds14 | 14.32 |
| Variable surface lipoprotein Y | MAG7080 | 11.55 |
| Variable surface lipoprotein X | MAG7070 | 8.47 |
| Variable surface lipoprotein | MAG3600 | 7.49 |
| NADH oxidase | MAG2630 | 2.75 |
| Lipoprotein P48 | MAG0120 | 2.46 |
| Hexosephosphate transport protein | MAG4970 | 2.34 |
| Lipoprotein | MAG6200 | 2.17 |
| Small ribosomal subunit protein uS3 | MAG5400 | 1.70 |
| Lipoprotein P80 | MAG5030 | 1.49 |

<sup>a</sup> Proteomic analysis of the anti-CDS14 and anti-P48 eluates
following co-incubation of PG2<sup>T</sup>[ICEA]<sup>G</sup> (ICEA donor) and pPG2<sup>P</sup>
(recipient); <sup>b</sup> Protein names according to UniProtKB/TrEMBL entries
in the UniProt Knowledgebase (UniProtKB;
https://www.uniprot.org/); <sup>b</sup> Accession number of the CDS from the
PG2 genome and ICEA in the GenBank database (accession number:
CU179680.1 and CT030003.1, respectively); <sup>c</sup> Intensity-Based
Absolute Quantification values based on a single .

**Table S3.**
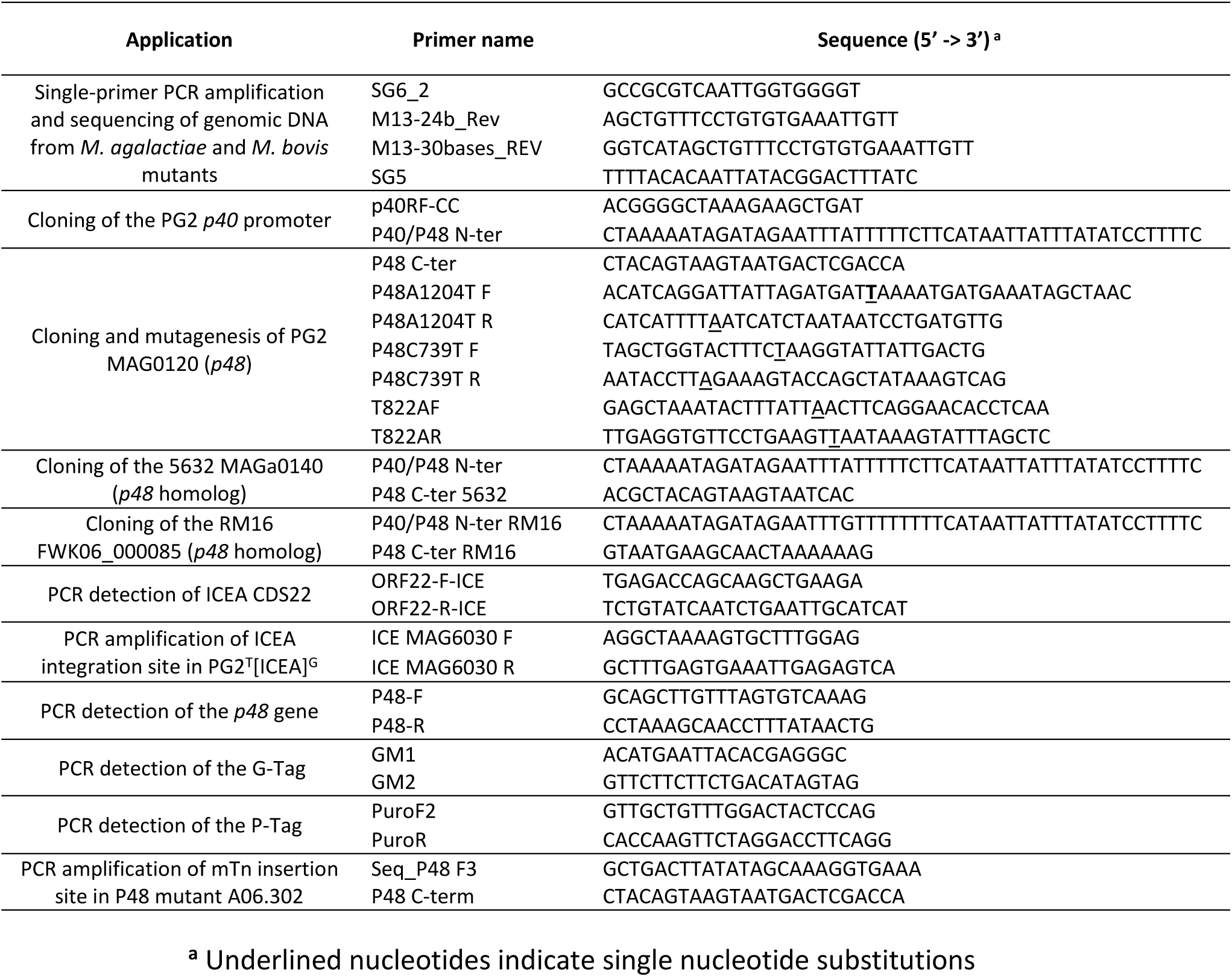
oligonucleotide primers.

## SUPPLEMENTARY FIGURES

**Figure S1.**
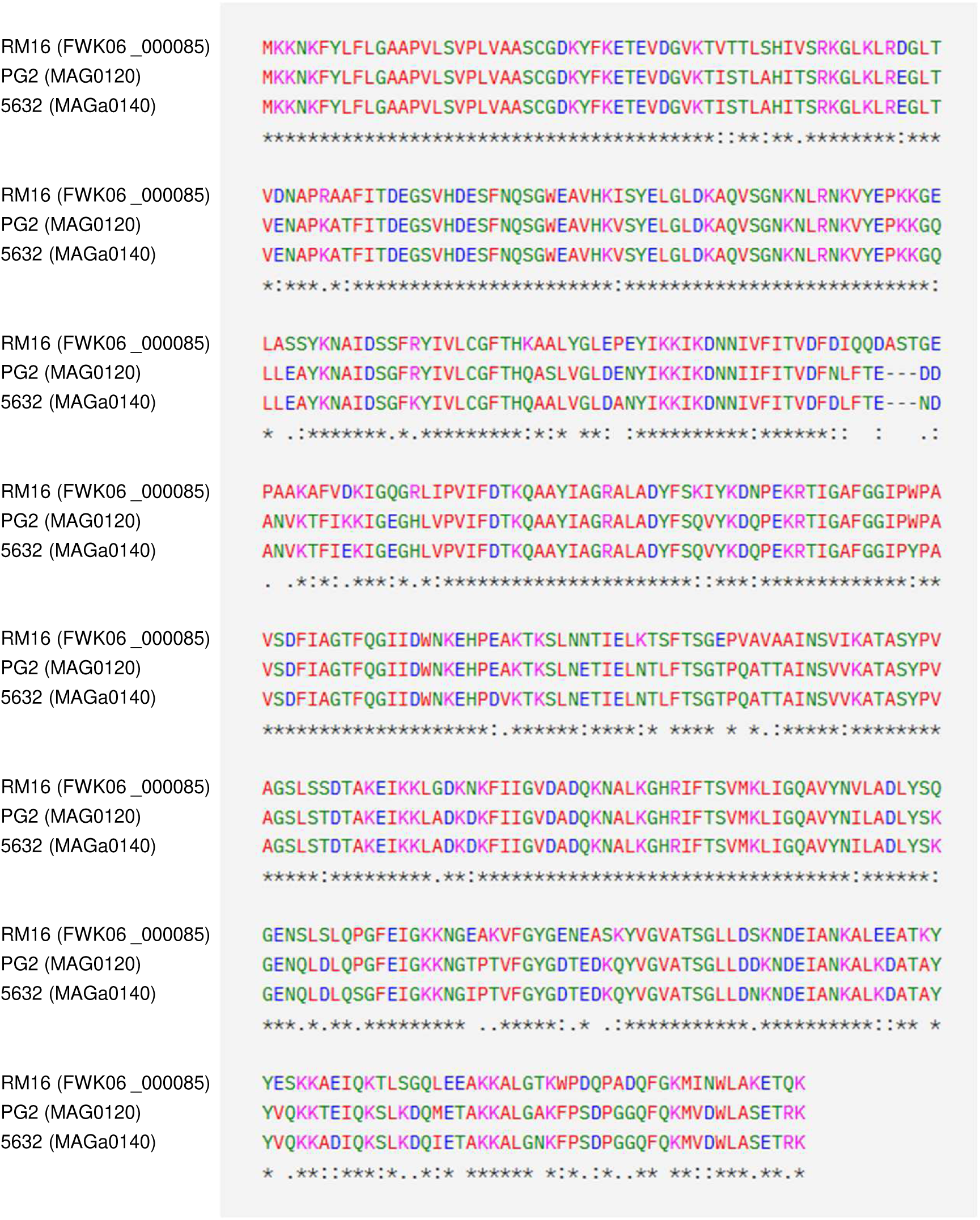
Multiple sequence alignment of the P48 lipoprotein from *M. agalactiae* PG2 and its homologs from *M. agalactiae* 5632 and *M. bovis* RM16 strains. Multiple sequence alignment by MUSCLE (3.8). CDS accession numbers in the GenBank database are indicated in parentheses.

**Figure S2.**
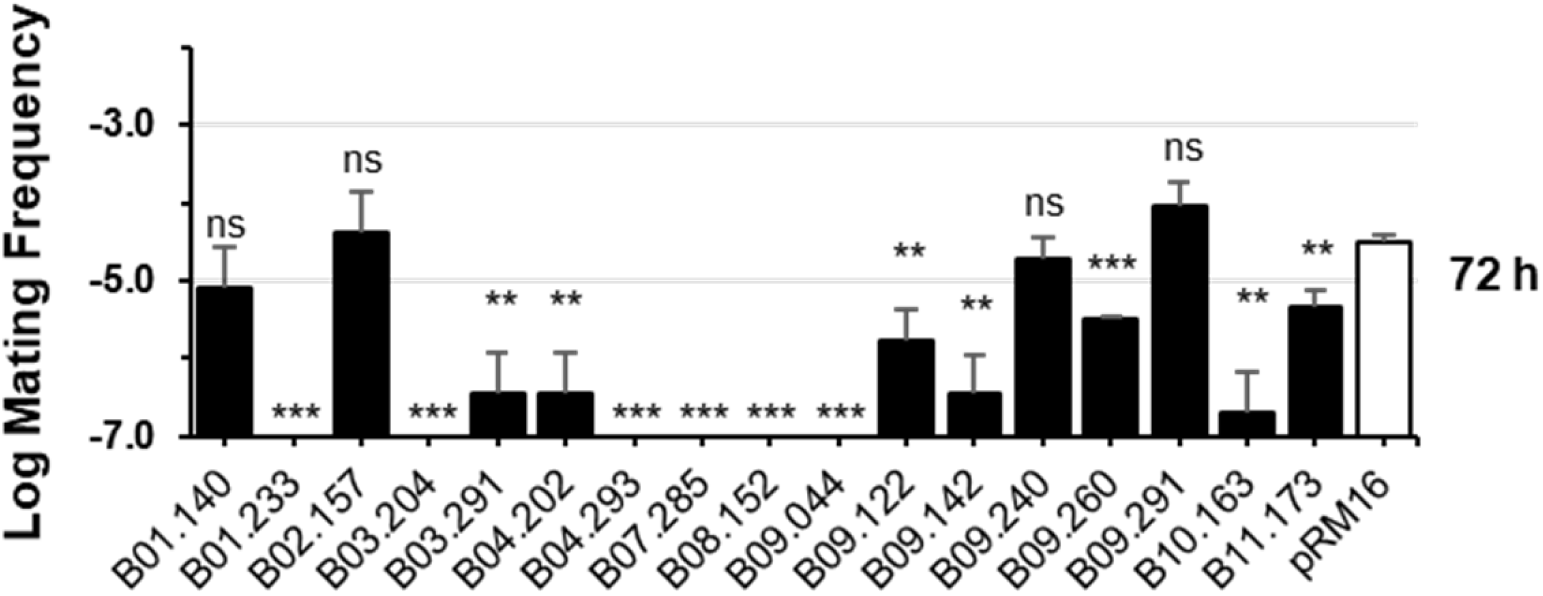
Resistance of *M. bovis* mutants to ICE transfer. ICE transfer from J228[ICEB]^G^ was measured to assess resistance. The recipient pRM16^P^ (pRM16) was used as positive controls (Table 1). Mating frequency was calculated as the ratio of dual-resistant transconjugants to total CFU. Data represent the mean ± SD of ≥3 independent experiments. Statistical significance was determined using two-tailed t-tests relative to pRM16^P^: ns, p ≥ 0.05; **, p < 0.01; ***, p < 0.001.

**Figure S3.**
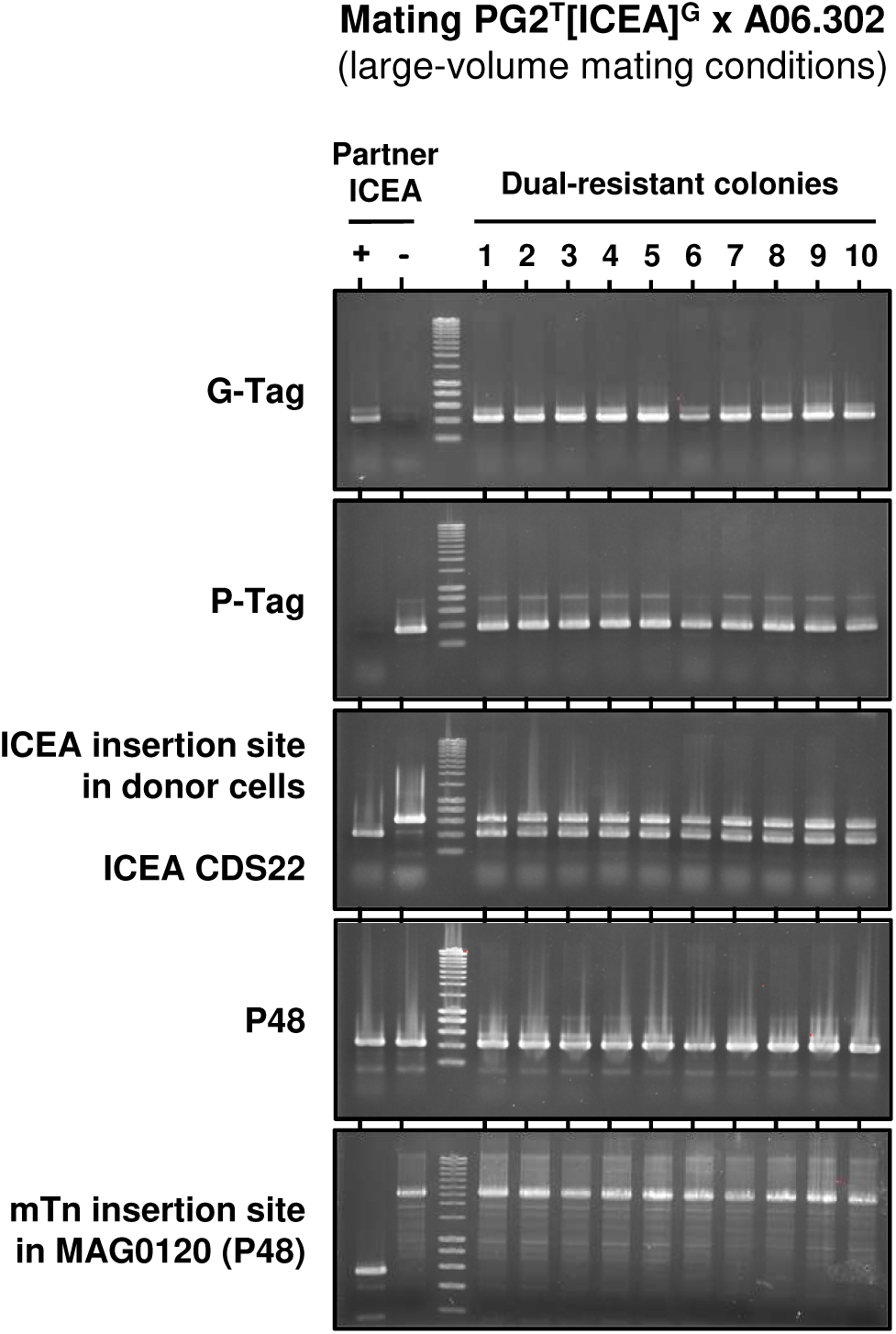
PCR characterization of dual-resistant colonies from mating PG2^T^[ICEA]^G^ x A06.302. PCR amplification of 10 dual-resistant colonies selected from large-volume mating conditions. All dual-resistant colonies were confirmed as transconjugants by PCR detection of antibiotic resistance markers originating from the ICEA donor PG2^T^[ICEA]^G^ (G-Tag) and from the ICEA recipient A06.302 (P-Tag). They were also tested positive for the ICEA CDS22 (ICEA CDS22) and the chromosomal region flanking the ICEA insertion site in the donor (ICEA insertion site in donor cells), indicating that in these transconjugants, the ICEA is integrated at a different chromosomal location. Additionally, the transconjugants were PCR positive for the *P48* gene MAG0120 (P48). However, PCR amplification of the region spanning the mTn insertion site in the P48 mutant A06.302 (mTn insertion site in MAG0120) yielded a large amplicon (> 2.5 Kb), confirming that the transconjugants are P48 mutants. The oligonucleotide primers used for PCR amplifications are described in Table S1. Mating partners were used as controls (ICEA+, PG2^T^[ICEA]^G^; ICEA-, A06.302).

**Figure S4.**
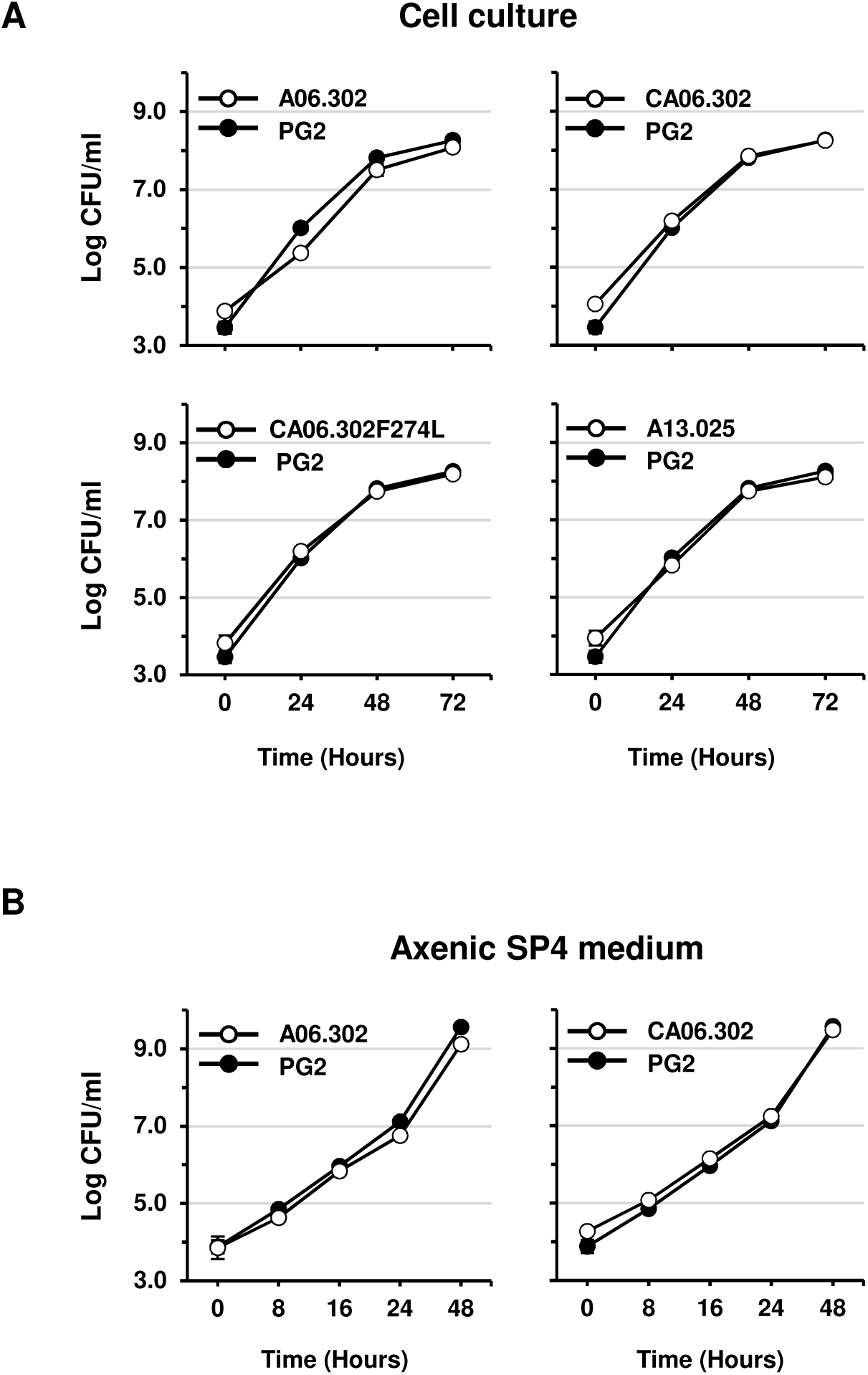
Growth curves of P48 mutants and complemented strains under cell culture and axenic conditions. Mycoplasma growth was monitored under (A) cell culture and (B) axenic conditions. Changes in CFU titers of wild-type PG2 (closed circles) were compared with those of the P48 mutant A06.302, the complemented strains CA06.302 and CA06.302F274, and the ABC transporter permease mutant A13.025. Data represent the mean of three independent experiments; error bars indicate standard deviations.

**Figure S5.**
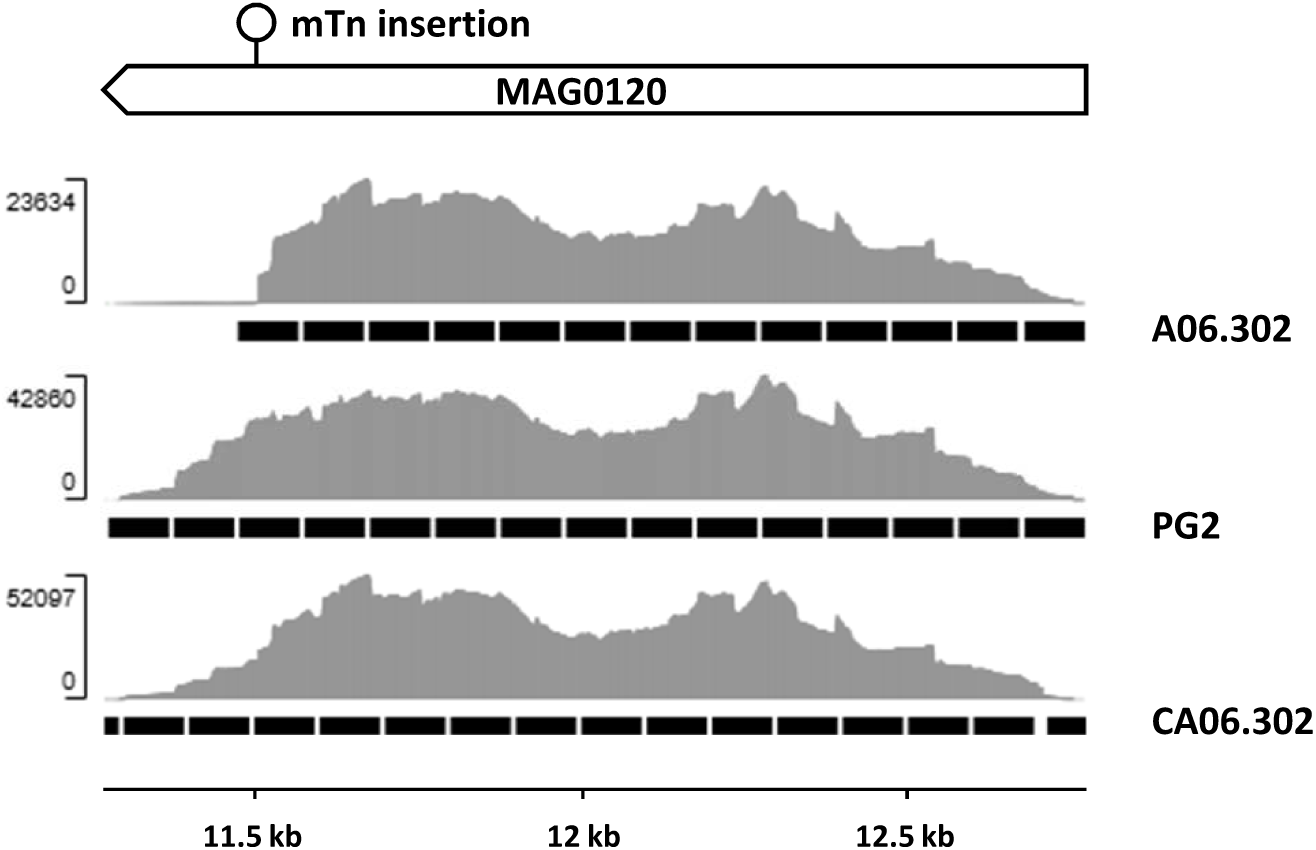
Detection of *p48* transcripts in the P48 mutant. RNA-seq detection of RNA transcripts from the *p48* gene (MAG0120) in the wild-type PG2 (PG2), the P48 mutant A06.302 (A06.302) and the complemented strain CA06.302 (CA06.302) generated by transformation of the P48 mutant A06.302 with plasmids pO/T-P48. The position of the transposon insertion site in MAG0120 is indicated (mTn). Y-axis shows read counts; black boxes denote gene regions with detected RNA transcripts.

**Figure S6.**
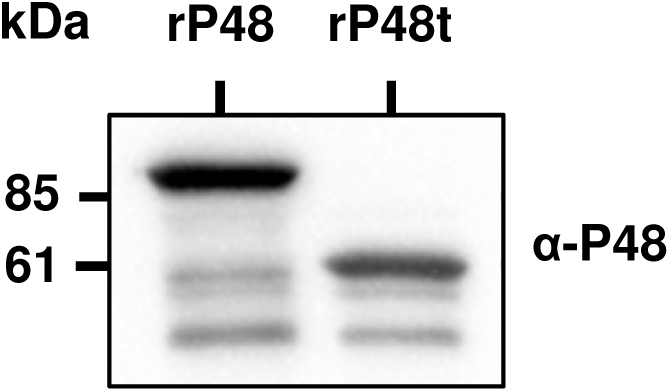
Reactivity of anti-P48 antibodies with a truncated recombinant P48 lipoprotein. Western blotting analysis of the reactivity of anti-P48 antibodies (α-P48) with full-length (rP48) and truncated forms of lipoprotein P48 (rP48t) expressed as recombinant soluble proteins in *E. coli.* The pH6HTN His₆-HaloTag® T7 vector (Promega) was used to express the soluble full-length (amino acid residues 26 to 465) and truncated (amino acid residues 26 to 246) P48 lipoprotein. Both proteins lack the N-terminal signal peptide (residues 1 to 25) and have an N-terminal His₆-HaloTag of approximately 35 kDa. Western blotting experiments were carried out using total protein extracts from *E. coli*.

**Figure S7.**
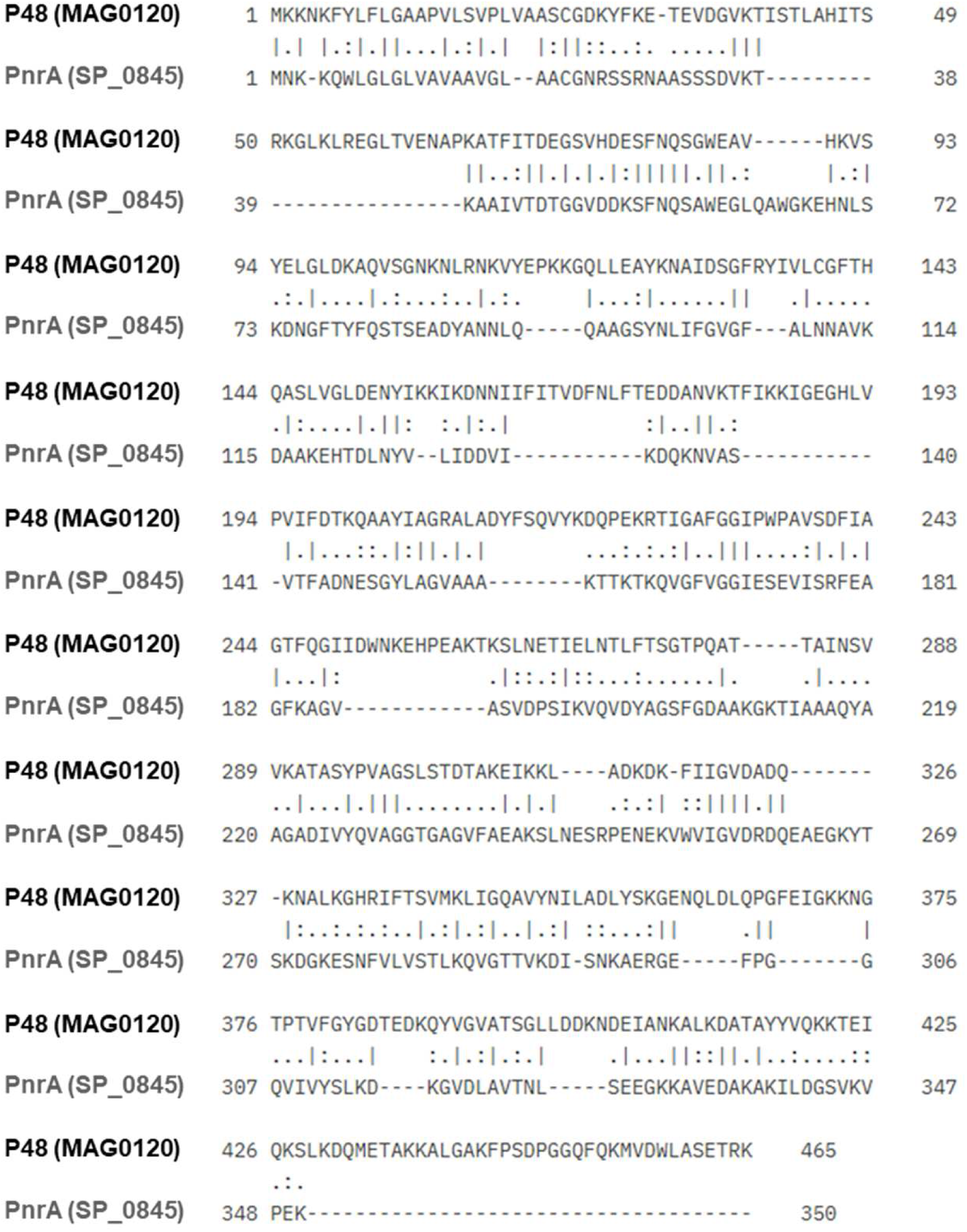
Sequence alignment of the *M. agalactiae* P48 and *S. pneumoniae* PnrA. Global sequence alignment of *Mycoplasma agalactiae* PG2 lipoprotein P48 (P48) and *Streptococcus pneumoniae* TIGR4 PnrA (PnrA) using EMBOSS Needle (https://www.ebi.ac.uk/jdispatcher/psa/emboss_needle). The *p48* and *pnrA* gene access numbers are indicated in parentheses.

